# EpiSegMix: A Flexible Distribution Hidden Markov Model with Duration Modeling for Chromatin State Discovery

**DOI:** 10.1101/2023.09.07.556549

**Authors:** Johanna Elena Schmitz, Nihit Aggarwal, Lukas Laufer, Jörn Walter, Abdulrahman Salhab, Sven Rahmann

## Abstract

**Motivation:** Automated chromatin segmentation based on ChIP-seq data reveals insights into the epigenetic regulation of chromatin accessibility. Existing segmentation methods are constrained by simplifying modeling assumptions, which may have a negative impact on the segmentation quality.

**Results:** We introduce EpiSegMix, a novel segmentation method based on a hidden Markov model with flexible read count distribution types and state duration modeling, allowing for a more flexible modeling of both histone signals and segment lengths. In a comparison with two existing tools, ChromHMM, Segway and EpiCSeg, we show that EpiSegMix is more predictive of cell biology, such as gene expression. Its flexible framework enables it to fit an accurate probabilistic model, which has the potential to increase the biological interpretability of chromatin states.

**Availability and implementation:** Source code: https://gitlab.com/rahmannlab/episegmix.

## 1. Introduction

Each cell in a eukaryotic organism contains the same genetic information to build all required structural and functional gene products. However, cell-to-cell variation is essential for having specialized tissues with distinct physiological functions and to adapt to environmental changes (Cavalli and Heard, 2019; Carter and Zhao, 2021). This necessitates an additional layer of processes regulating gene expression to enable cell differentiation and to maintain cellular identities throughout cell divisions (Allis and Jenuwein, 2016). Among the mechanisms tightly regulating gene expression are transcription factors and epigenetic modifications, like DNA methylation and histone modifications.

### Histone modifications

With increasing knowledge about the role of histone modifications in altering the chromatin structure and DNA accessibility, it became apparent that different histone modifications are enriched in chromatin regions with distinct functional roles (Baker, 2011). For example, modification H3K4me3 (in histone protein H3, the lysine at position 4 (K4) is trimethylated (me3)) is enriched in promoters and can be linked to transcriptional activation; H3K36me3 is enriched in active genes, and H3K27me3 can be associated with gene repression by the Polycomb protein complex (Blackledge and Klose, 2021). Densely packed chromatin, called heterochromatin, is typically characterized by low levels of acetylation, whereas open, actively transcribed chromatin, called euchromatin, shows enrichment of acetylated lysine (Bannister and Kouzarides, 2011). Combinatorial patterns of multiple histone modifications allow us to characterize so-called *chromatin states* that describe the different functional states of both coding and non-coding regions in the genome (Baker, 2011).

### ChIP-seq

Advances in chromatin immunoprecipitation followed by sequencing (ChIP-seq) boosted the understanding of the various roles of histone modifications by enabling the generation of genome-wide histone maps in high-throughput experiments (Barski et al., 2007). For ChIP-seq, DNA-binding proteins of interest, such as specifically modified histones or transcription factors, are tagged with specific antibodies. After chromatin shearing, DNA fragments bound to the desired proteins or protein-modifications are captured and the bound DNA is extracted for sequencing. Reads are mapped to the reference genome to infer the positioning of histone marks across the genome (Park, 2009). The positional enrichment of reads is compromised by a certain amount of noise.

### Probabilistic models for segmentation

The availability of genome-wide ChIP-seq data led to the development of automated methods for genome segmentation and annotation. These methods use a probabilistic model to detect recurrent patterns of epigenetic marks using the aligned reads to determine the signal intensity at different positions in the genome (Fig. 1). HMMSeg (Day et al., 2007), ChromHMM (Ernst and Kellis, 2010) and EpiCSeg (Mammana and Chung, 2015) are based on hidden Markov models (HMM), Segway and Segway 2.0 (Chan et al., 2018) fit a Gaussian mixture model using a dynamic Bayesian network, and Daneshpajouh et al. (2022) developed a state–space model assuming that the observed data is a linear function of the state-specific parameter matrix plus Gaussian noise. For a more comprehensive review, we refer to Libbrecht et al. (2021).

**Figure 1.**
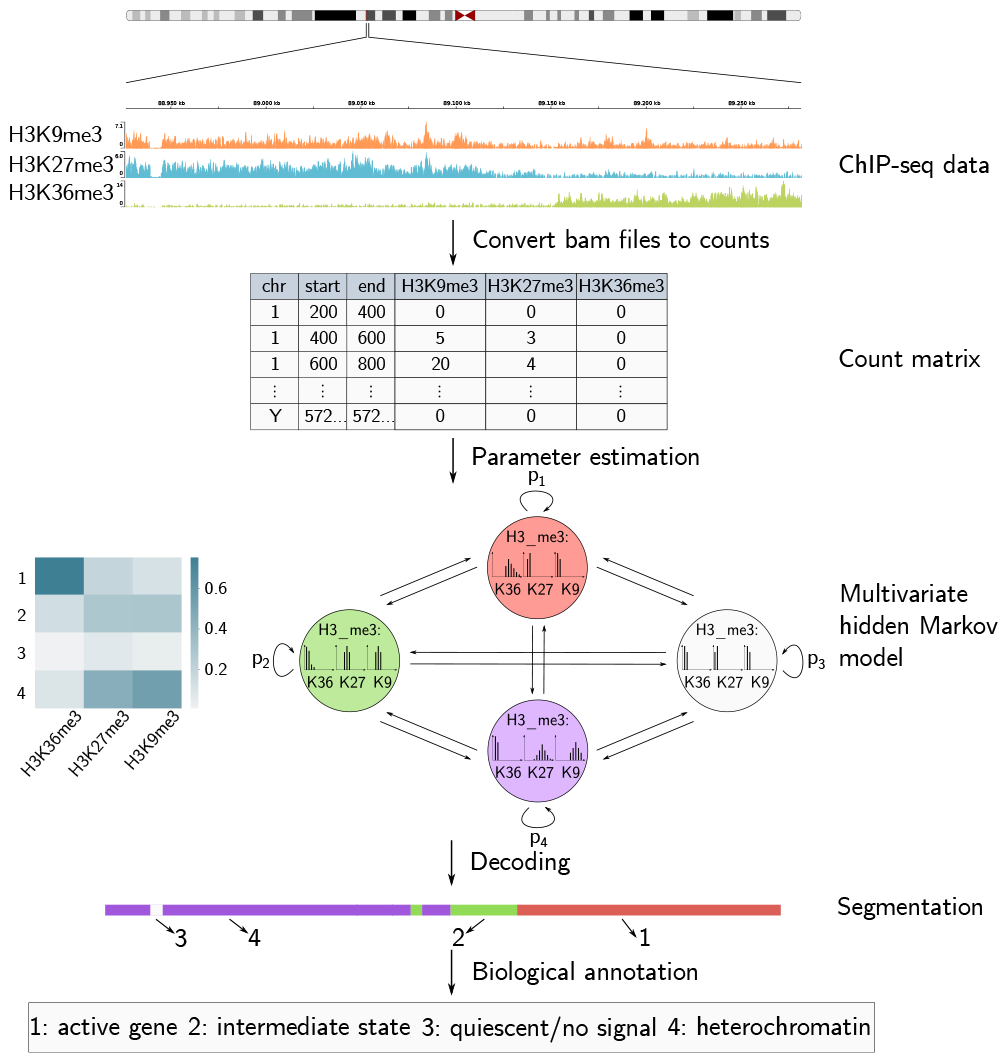
Chromatin segmentation: The reads of a ChIP-seq experiment are converted into a count matrix by counting the number of reads mapping to each non-overlapping 200 bp genomic interval. The states of a multivariate Hidden Markov model (HMM) capture patterns in the multivariate read count distribution of the histone marks, and the transition probabilities between states capture the relations between adjacent chromatin states.

### HMMs

Multivariate HMMs capture both combinatorial patterns of multiple histone marks and adjacency relations between different genomic elements, which makes them a prominent tool for chromatin state discovery (Lee and Park, 2014). An HMM describes two stochastic processes, an invisible Markov chain consisting of a finite set of hidden states and a visible process of observable signals. Here, the hidden states correspond to chromatin states and the observable signals to the observed read count vector per genomic region.

Each hidden state has state-specific probabilities of emitting an observation, called *emission probabilities*. For chromatin segmentation, the genome is divided into non-overlapping intervals (of typical length 200 bp), such that each observation is a vector of counts corresponding to the number of reads assigned to the interval per histone mark. Thus, the emission probabilities of a single hidden state capture a specific combinatorial pattern of multiple histone marks. Different states may hence define different functional genomic elements, such as promoters, enhancers or gene bodies. In addition, the relations between adjacent chromatin states are modeled via transition probabilities, which determine, for each state, the probabilities to either stay in the same state or to transition to another state.

### Modeling assumptions

The ability of an HMM to detect patterns that correspond to biologically meaningful chromatin states is constrained by the modeling assumptions underlying the emission and transition probabilities. These assumptions are thus a distinguishing feature of existing HMM-based methods. For example, ChromHMM fits an HMM on binarized data (high vs. low read count), where the emission probabilities are assumed to be independent Bernoulli experiments, and EpiCSeg models the emission probabilities using a Negative Multinomial distribution. Previous analyses of ChIP-seq data have shown that the read count distributions in some states and for some histone marks may be overdispersed and skewed, partly caused by differential protection against sonication, unequal binding affinity of distinct antibodies, sequence dependent PCR amplification and discrepancies when mapping to repeat-rich regions, which all introduce bias to the data (Diaz et al., 2012). Not all observed combinations of overdispersion and skewness can be captured by the commonly used probability distribution families, such as the Negative Binomial distribution. Furthermore, Beacon et al. (2021) showed that histone marks that are enriched in short domains, like promoters or TSS, are typically characterized by narrow peaks with high signal intensities, while histone marks enriched in broad domains, like heterochromatic regions, have lower signal intensities. Thus, regions in the genome covered by the same chromatin state may have different lengths, for example, short promoters and long heterochromatic regions. Existing tools only model a single duration distribution type (Geometric) with exponentially decreasing probabilities, and can only fit the mean length of a region, but not its shape to the observed data.

### Novel contributions

We propose a new flexible HMM architecture that relaxes the modeling assumptions of existing HMM-based methods in two ways. First, we allow to choose, for each histone modification, a different discrete distribution type from a broad selection (Table 1). This allows us to model more flexible read count distribution shapes, including overdispersed and skewed distributions. Second, we provide flexible duration modeling (using a state extension technique) to capture the characteristics of broad and narrow chromatin domains. By applying our method to publicly available ChIP-seq data, we show that such a flexible HMM leads to a better model fit and may increase segmentation accuracy and biological interpretability.

**Table 1:** Overview of available discrete distributions (Johnson et al., 1993).

| Name | Parameters |
| --- | --- |
| Binomial | $n \in \mathbb{N}, p \in [0, 1]$ |
| Poisson | $\lambda \in \mathbb{R}^+$ |
| Negative Binomial | $r \in \mathbb{R}^+, p \in [0, 1]$ |
| Beta Binomial | $n \in \mathbb{N}, \alpha \in \mathbb{R}^+, \beta \in \mathbb{R}^+$ |
| Beta Negative Binomial | $r \in \mathbb{R}^+, \alpha \in \mathbb{R}^+, \beta \in \mathbb{R}^+$ |
| Sichel | $\mu \in \mathbb{R}^+, \sigma \in \mathbb{R}^+, v \in \mathbb{R}$ |

## 2. Methods

### 2.1. Probabilistic Model

A hidden Markov model (HMM) is formally defined as a quintuple (*N, π*, Σ, *A, B*), where { 1, 2, …, *N*} is a finite set of hidden states, *π* is a probability vector with the starting probabilities for each state, Σ is a set of observable emission values, *A* is an *N* × *N* matrix, where each entry *a*_*ij*_ denotes the transition probability to move from state *i* to state *j*, and *B* = (*b*_*j*_(·)| *j* ∈ {1, 2, …, *N*}) is the complete collection of parameters required to calculate the emission probabilities of an observation in each state *j* (Lee and Park, 2014). For a sample with *T* observations or time points, the collection of random variables is thus given by (*O, Q*), where *O* = (*O*_1_, …, *O*_*T*_) denotes the observed sequence and *Q* = (*Q*_1_, …, *Q*_*T*_) denotes the hidden state sequence.

Due to the HMM independence assumptions (*Q* is a Markov chain, *O*_*t*_ is conditionally independent of everything else, given *Q*_*t*_; Bilmes (1998)), the probability that an HMM with parameters *θ* = (*π, A, B*) generates the combination (*Q* = *q, O* = *o*) is

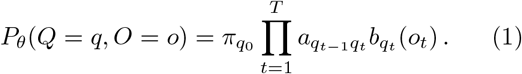

For given fixed *O* = *o*, the Viterbi algorithm (Rabiner, 1989) determines the state sequence *q* that maximizes this probability (for fixed transition and emission probabilities).

For chromatin segmentation, the emission alphabet is given by the multivariate, countable infinite set

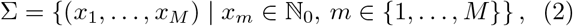

where *M* is the dimension of the emission alphabet, corresponding to the number of histone modifications in the input data. Hence, the natural choice is to model the emission probabilities using a multivariate discrete distribution. Under the assumption that the read counts of all histone modifications are conditionally independent of each other given a state, the emission probability for an observation *o*_*t*_ = (*o*_*t*1_, …, *o*_*tM*_) in state *j* is given by

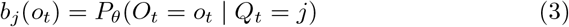

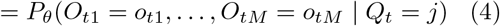

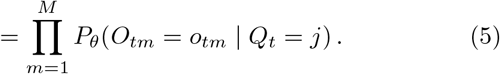

This leads to a flexible framework in which different distribution types may be selected to model the read counts of distinct histone marks. Table 1 gives an overview over the univariate discrete distributions available in EpiSegMix (see Supplementary Section S1 for details). Distributions with more parameters are more flexible and hence lead to a more accurate model fit: One-parameter distributions (Poisson) may fit the mean of the observed read counts, but not variance or skewness; two-parameter distributions (e.g., Negative Binomial) may fit both mean and variance but not skewness; three-parameter distributions may fit all three moments. With data from several thousand genomic intervals per state, there is no danger of over-fitting three parameters. Still, each distribution has its own limitations and dependencies between moments, so having a variety of options is beneficial. The best performing distribution may vary for histone modifications and between experiments with different data quality. Therefore, we provide a workflow to find for each mark the distribution that maximizes the log-likelihood. In general, the Sichel and Beta Negative Binomial distribution are well suited for modeling histone counts due to their high flexibility. For further information, see Supplementary Section S4.

### 2.2. Duration Modeling

The typical HMM topology is a fully connected graph, including self-loops on states. Hence the sojourn time *X* in a state follows a Geometric distribution *P* (*X* = *k*) = *p*· (1 −*p*)^*k*−1^ for some *p >* 0, with exponentially decreasing probabilities for longer durations (Rabiner, 1989). Although the mean of a Geometric distribution can be made arbitrarily large, the variance increases with it and the stays at *P* (*X* = 1). Geometric distributions model durations that are short with higher probability, but have limited flexibility when modeling longer durations with a mode far away from 1. We therefore propose to use an extended-state HMM architecture, in which each state is internally represented as a sub-HMM. Each sub-HMM has a linear left-to-right topology with a different number of sub-states, but with the additional constraints that all sub-states have the same emission probabilities and same self-transition probabilities (Russell and Cook, 1987); see Fig. 2A. With this topology, the state duration follows a Negative Binomial distribution. It has two parameters: the state-exit probability *p* (same as before), and the copy number *r* of the state (*r* = 1 gives the Geometric distribution). In contrast to a Geometric distribution, the mean and variance of a Negative Binomial distribution can be controlled independently, and the mode can be placed at an arbitrary duration (Fig. 2B). Therefore, both long and short durations can be modeled flexibly.

**Figure 2.**
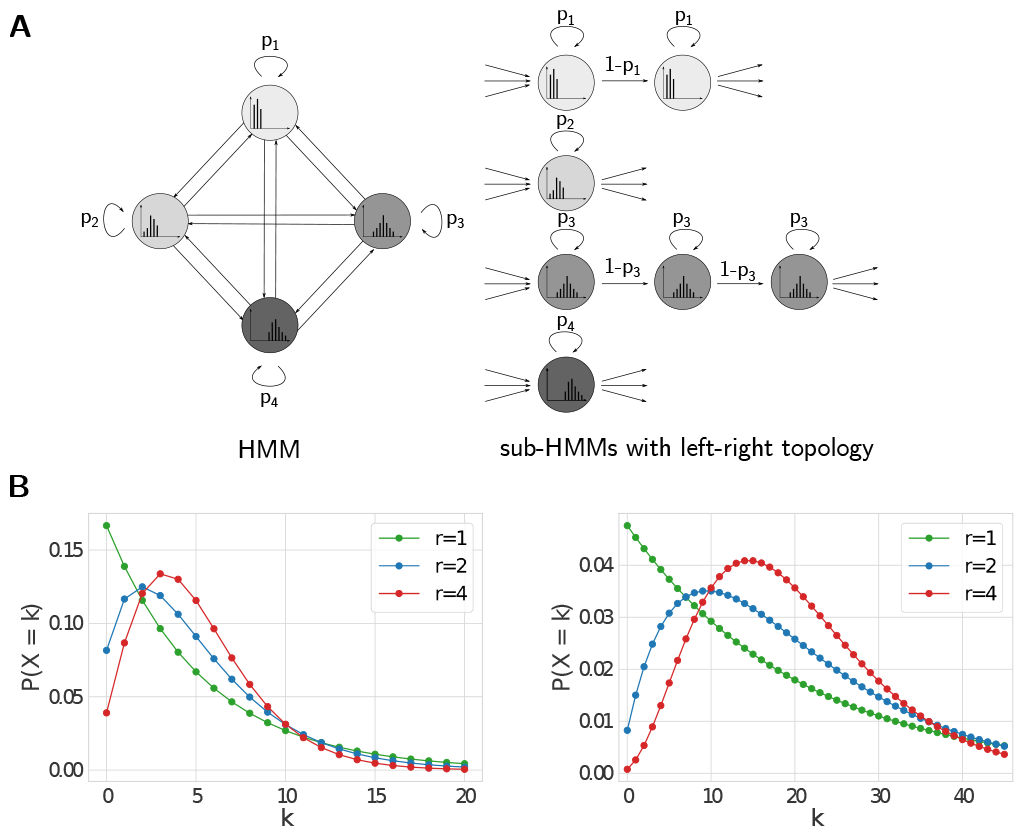
Duration modeling. **A** Extended-state HMM with different numbers of states in the sub-HMMs for a univariate 4-state HMM. **B** Comparison of state duration distribution for a state in a classical HMM (*r* = 1) and in a topology HMM with 2 or 4 sub-states to achieve the same mean of 5 (left) or 20 (right).

To determine a suitable number of states for each sub-HMM, the segment lengths are computed for each state under the current model. The number of substates are then given by the estimated parameter *r* of a Negative Binomial distribution under the additional constraint that *r* is a natural number between one and five. By default, the topology is adjusted twice. Between the adjustments, the parameters of the HMM are estimated for a fixed number of iterations (default 5) and for the final topology until convergence.

### 2.3. Parameter Estimation

Before an HMM can be used to estimate the state sequence, its parameters (transition and emission probabilities) must be estimated. As we typically do not have labeled training data, parameter estimation must proceed in an unsupervised manner using the Baum-Welch algorithm, which is a concretization of the general expectation maximization (EM) algorithm tailored to the structure of HMMs (Rabiner, 1989). It alternates between estimating the model parameters (M-step), given a (fuzzy or probabilistic) assignment of observations to states, and re-estimating the state membership probabilities of each observation (E-step) until convergence (details in Supplementary Section S1).

## 3. Implementation

The flexible distribution HMM and parameter estimation is implemented in C++ as a command-line tool EpiSegMix. Its source code is at https://gitlab.com/rahmannlab/episegmix.

All steps of the surrounding workflow are incorporated in a `Snakemake` workflow (Mölder et al., 2021).

By default, we estimate parameters on the ENCODE pilot regions which contain a good representation of the whole genome and are thus commonly used to fit the model (Hoffman et al., 2012; Daneshpajouh et al., 2022). Alternatively, the user may specify a list of chromosomes to be used for model fitting. The segmentation is performed genome-wide.

The main output of chromatin segmentation is a file that assigns one state to each position in the genome, and an HTML report with plots that characterize the model and segmentation, enabling their biological interpretation. For example, the heatmap showing the normalized histone modification intensities of each state (as in Fig. 6B) is central to determine the genomic function of the states.

## 4. Results

We evaluated EpiSegMix on publicly available ChIP-seq data for the human cell lines K562, HepG2, GM12878, IMR90, H1 and SJCRH30 provided by the ENCODE consortium (Dunham et al., 2012) using the most recently processed data. With the selected cell lines, we evaluate our method on ChIP-seq experiments that were performed over the last decade (from 2010 to 2020) with different data properties (e.g., the mapped read length ranges from 36 bp to 100 bp). To analyze the robustness of all methods, we generated the count matrix for two replicate experiments each (see Supplementary Section S6). To reproduce the results, all accession numbers are provided in Supplementary Section S9 and the script to create the count matrix is part of the code repository. We restricted our analysis to the six core histone marks H3K9me3, H3K27me3, H3K36me3, H3K4me1, H3K4me3 and H3K27ac defined by the IHEC consortium (Bujold et al., 2016).

Since chromatin segmentation is an unsupervised method and no ground truth is available, we evaluate the performance of EpiSegMix by comparing it to three established chromatin segmentation tools ChromHMM (Ernst and Kellis, 2012), EpiCSeg (Mammana and Chung, 2015) and Segway (Chan et al., 2018).

We first perform a quantitative comparison by evaluating how well the different methods can predict gene expression and ATAC-seq data. Afterwards, we analyze how the different methods reflect known genome biology by showing further characteristic plots for one exemplary dataset.

### 4.1. Data Processing

In a preprocessing step, we convert the aligned reads into a count matrix using the *bamsignals* package (Mammana and Helmuth, 2023). In the count matrix, each row corresponds to a consecutive, non-overlapping region with a fixed window size (default 200 base pairs), called bins, and each column corresponds to a distinct histone mark. Each read is assigned to exactly one genomic bin depending on the position of its 5’ end.

For all methods, we fitted a 10-state model. EpiSegMix and Segway were trained on the ENCODE pilot regions of *hg38*. For EpiSegMix, we determined the best-suited distributions for each dataset based on the fit for each mark individually, listed in Supplementary Section 4. For Segway, the *resolution* was set to 200 bp, the *track-weight* to 0.01 and the *segtransition-weight-scale* and *prior-strength* to 1. For ChromHMM and EpiCSeg, the default parameters were used (200 bp resolution, 10-state model).

### 4.2. Advantages of Flexible Distribution Modeling

To show the advantages of flexible emission and duration distribution types for chromatin segmentation, we compare fitted models using different emission distributions and with and without duration modeling. Narrow marks, such as H3K4me3, often have skewed read count distributions. Fitting the Negative Binomial (2 parameters) and Sichel (3 parameters) distribution to the read counts of H3K4me3 shows the limitation of the Negative Binomial distributions to model highly skewed and overdispersed data (Figure 3A). Further evaluation of the effect that emission modeling has on the segmentation is provided in Supplementary Section S4.

**Figure 3.**
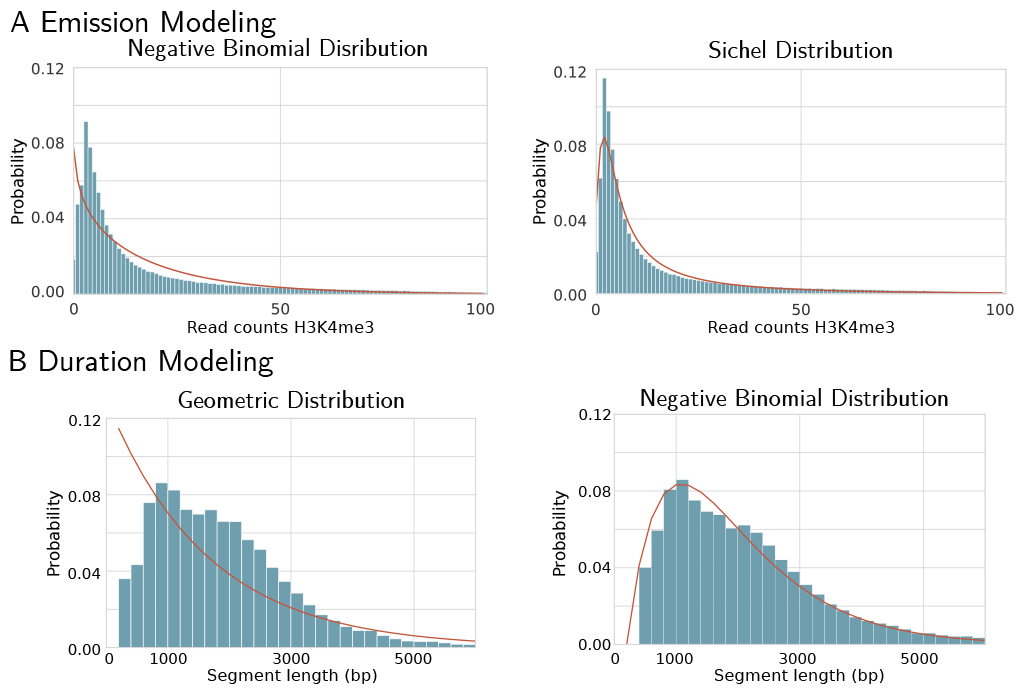
Flexible emission and duration modeling. The histograms show the sample distribution and the red curves the theoretical distribution fitted by the model. **A** Results of fitting a 3-state HMM to the mark H3K4me3 using the Negative Binomial and Sichel distribution (for the state with high H3K4me3 in HepG2 1). **B** State duration in the HepG2 1 promoter state for an HMM with a classic (Geometric) and extended (Negative Binomial) topology.

Figure 3B shows that the state duration, determining the segment length (number of consecutive bins assigned to the same state), does not follow a Geometric distribution for most chromatin states. In comparison, the Negative Binomial distribution, as fitted by our flexible duration model, leads to a more accurate description of the real segment length distribution.

### 4.3. Evaluation of Gene Expression Prediction

Since a biologically meaningful segmentation should have states that correlate with different gene expression levels, we compared how well the chromatin states of EpiSegMix, EpiCSeg, ChromHMM and Segway can predict gene expression. To measure the gene expression in each 200 bp bin that contains (part of) a protein-coding gene, we used total RNA-seq experiments for the different cell lines provided by ENCODE and assigned each bin the log(FPKM + 1) normalized expression value of the gene (FPKM: fragments per kilobase of transcript per million mapped reads). We performed linear regression with the state labels as categorical predictors, i.e., for each bin *i*’s true expression *x*_*i*_, we used the state-specific mean expression as predictor 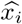 and measured the mean quadratic regression error vs. the mean quadratic error using the global mean 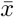 as predictor and computed the coefficient of determination 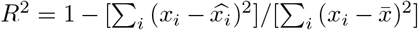 (between 0 and 1, where 1 is perfect). The *R*^2^ values vary for the different cell lines, which can partly be explained by the unequal data quality. Fig. 4 compares the *R*^2^ values of the different methods across cell lines. On average, EpiSegMix has the highest predictive power among the methods.

**Figure 4.**
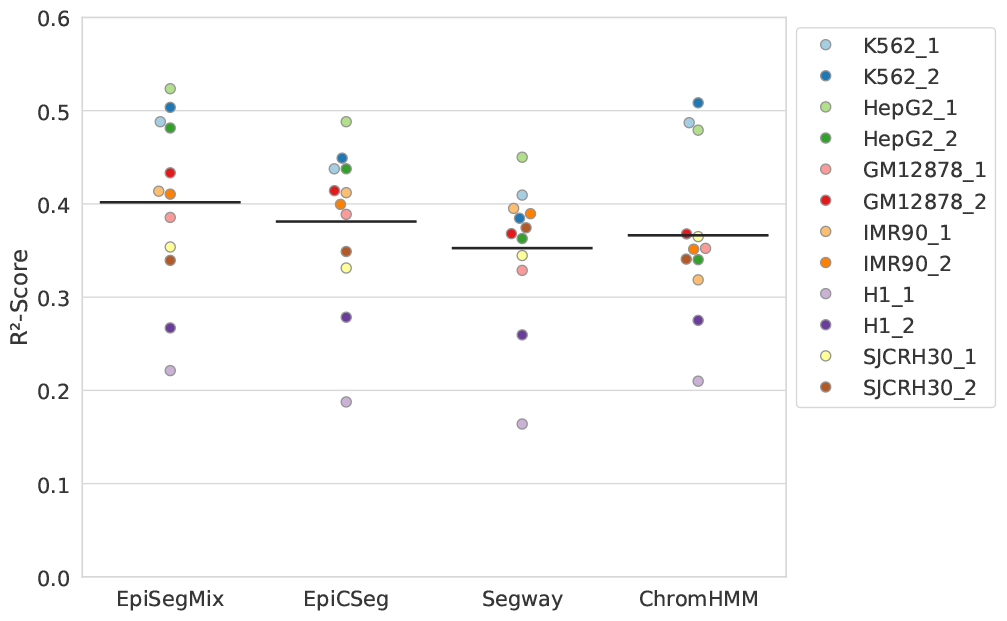
Prediction of transcription levels from chromatin state labels: coefficient of determination *R*^2^ (y-axis) using state labels as categorical predictors for log(FPKM+ 1) expression values for the different methods (x-axis) on different cell lines (color). Black lines indicate mean *R*^2^ for each method.

### 4.4. Evaluation of ATAC-seq Prediction

We performed a similar analysis predicting ATAC-seq read counts instead of gene expression levels for all cell lines with available ATAC-seq experiments. We counted how many reads of the ATAC-seq experiment map to each non-overlapping 200 bp bin in the genome to generate a count matrix in the same way as for the histone counts. In the same way as for gene expression, we performed linear regression to predict log-transformed ATAC-seq read counts using the state labels as categorical predictors. Since ATAC-seq measures the chromatin accessibility, active states, like promoter or transcription states, should be predictive of high ATAC-seq counts, while heterochromatic states should be predictive of low ATAC-seq counts. Fig. 5 shows that EpiSegMix and EpiCSeg have a similar predictive power of chromatin accessibility, as measured by the coefficient of determination *R*^2^, while ChromHMM and Segway have lower *R*^2^ scores on average.

**Figure 5.**
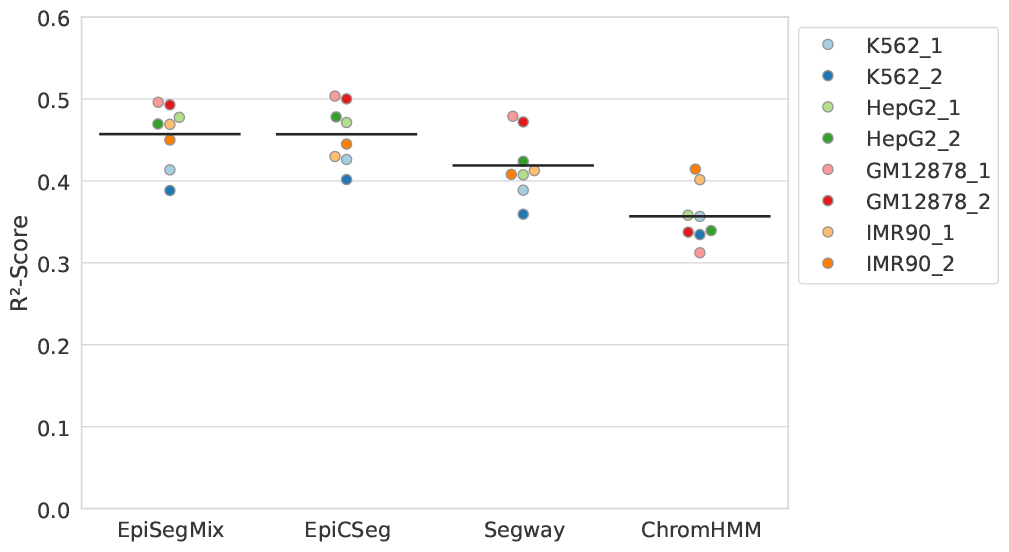
Prediction of ATAC-seq levels from chromatin state labels. The plot shows the coefficient of variation (*R*^2^) using standard linear regression with the state labels as categorical predictors to predict log transformed ATAC counts. The black line shows the mean *R*^2^ value.

**Figure 6.**
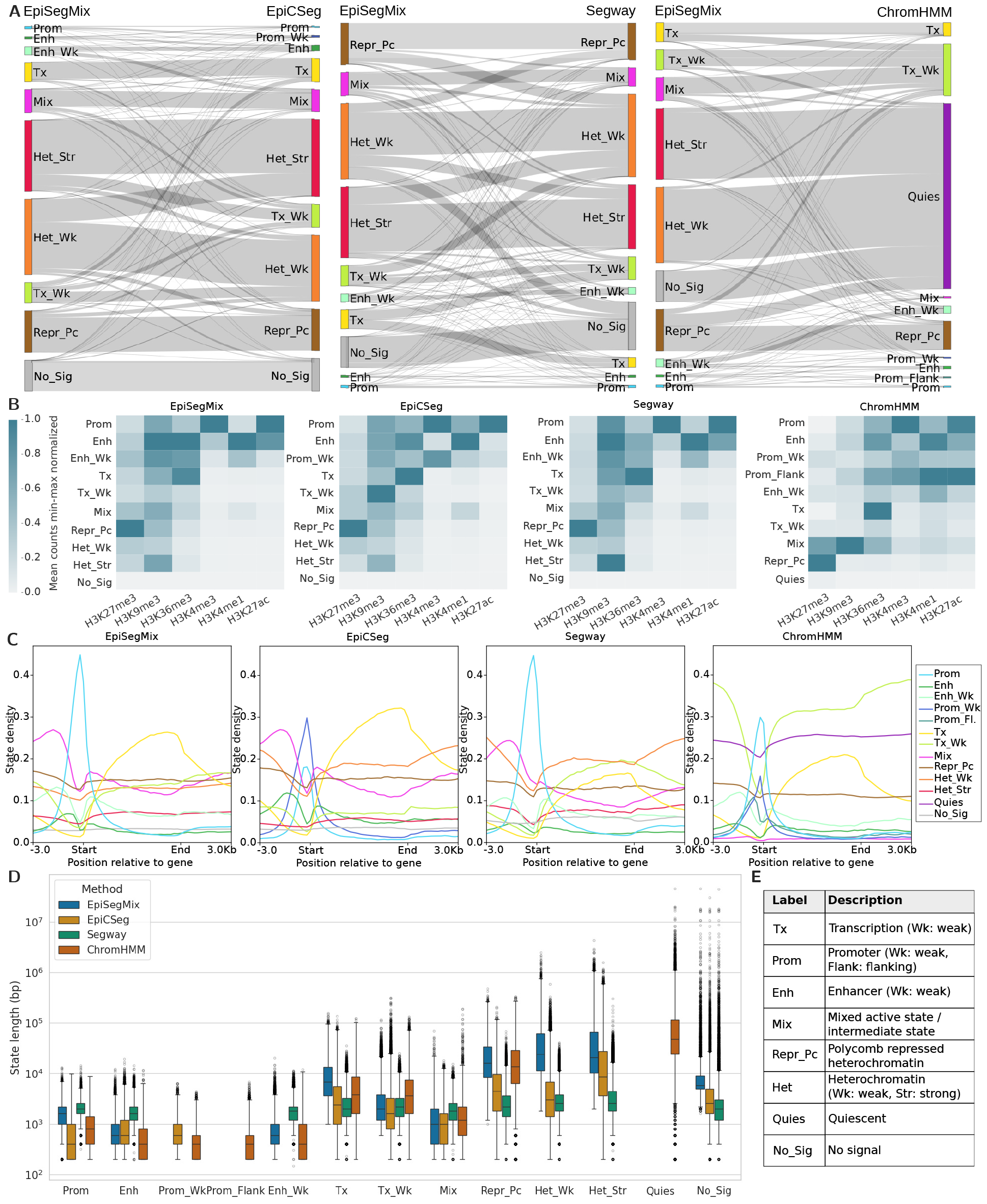
Comparison between EpiSegMix and EpiCSeg, Segway and ChromHMM for K562 1. **A** State overlap between methods. The bar heights corresponds to the genomic coverage of the biologically annotated state and the edge thickness to the overlap between the methods. **B** Heatmaps showing the histone enrichment per state. The mean counts per state are normalized column wise, such that for each histone modification the state with the maximum mean count has a value of one and the state with the lowest mean count has a value of zero. **C** State distribution. The line plots show how often each state occurs around protein coding genes. **D** State duration. The plot shows for each biologically annotated state the state length distribution in base pairs of the different methods. **E** State description.

### 4.5. Evaluation of Reflected Genome Biology

We perform a qualitative analysis of the different methods by comparing the similarities of the genome-wide segmentation and functional assignment of states, their genome coverage, overall segment length and enrichment around protein coding genes (Fig. 6). An in-depth example (genome browser view) is provided in Supplementary Section S7. For better comparability, we manually assigned a label that best describes the biological function of the state to each numerical state ID. This facilitates the comparison of state assignments between methods. A description of each label is given in Fig. 6E and Supplementary Section S8.

Fig. 6A shows that the states assigned by the count based methods EpiSegMix, EpiCSeg and Segway have a similar genome coverage and high overlap between state assignments. Moreover, all three methods EpiSegMix, EpiCSeg and Segway provide a deeper epigenomic distinction of heterochromatic regions that ChromHMM aggregates into a single state. EpiSegMix discriminates a “no signal” state from a strong and a weak heterochromatic state that show distinctive patterns of enriched histone marks. Thereby, EpiSegMix better captures transitions from closed to open chromatin as compared to ChromHMM (Supplementary Section S7). This distinction is also supported by Segway and EpiCSeg. In addition, EpiSegMix defines a more consistent and accurate label for unmappable regions in comparison to all other methods (Supplementary Section S6). While, in a 10 state model, ChromHMM appears to provide a more fine-grained distinction of regulatory states, such as weak and flanking promoter regions, the robustness of these classifications is not always given. Thus, a biological interpretation of these regions should be taken with caution (Supplementary Section S6).

Fig. 6B shows the genome wide enrichment of histone marks across states for each tool. All methods find similar patterns of histone marks: Enrichment of regulatory marks such as H3K27ac, H3K4me1 and H3K4me3, is predominantly observed in promoter and enhancer states, as expected. Actively transcribed mark H3K36me3 is enriched in transcription states; the repressive mark H3K27me3 is most strongly enriched in Polycomb repressed heterochromatin. Overall absolute levels of H3K9me3 are low, but due to the column-wise normalization in Fig. 6B, it appears enriched in a number of states, possibly reflecting overall background noise.

Despite the overall high state concordance across tools, some differences can be observed particularly in a gene centered comparison, i.e., across gene bodies, including ± 3 Kbp upstream and downstream of the gene transcription start and gene end, respectively (Fig. 6C). Coordinates were taken from ENCODE reference ENCFF824ZKD. EpiSegMix and Segway show a higher enrichment of promoter states around TSSs as compared to ChromHMM and EpiCSeg, which discriminate between weak and strong promoters. This distinction is not linked to a better prediction of transcription levels.

When comparing the segment lengths (i.e., consecutive genomic bins assigned to the same state) and their genomic distributions across the different methods we observe that the duration flexibility of EpiSegMix helps to capture the wide range of short (promoters/enhancers), intermediate (short to long genes) and long (heterochromatic regions) state durations in a more consistent manner as compared to all other methods. For example, EpiSegMix’ Tx state most effectively covers large genes (i.e., over 40% of all human genes are longer than 0.8 · 10^4^ bp), allowing for a more accurate annotation of genes among different classes. Another advantage of EpiSegMix in comparison to EpiCSeg and Segway can be observed in the longer segment lengths for Polycomb repressed genes (Repr Pc) and for heterochromatic regions (Het Wk and Het Str), which more closely match the known size of broad domains (Steensel and Belmont, 2017).

## 5. Discussion

We developed EpiSegMix, a flexible HMM framework for chromatin segmentation. We enhanced the flexibility of existing HMMs with respect to modeling both emission probabilities and state durations. To account for the overdispersed and skewed ChIP-seq read count distributions, the read counts of each histone modification can follow a different discrete distribution type. We currently support a variety of distributions, and further distributions may be added in the future. The internal HMM topology was adjusted to be able to model state durations that follow a Negative Binomial instead of a Geometric distribution, which better reflects the inherent segment length of chromatin states that cover either small peaks or broad domains. For example, lamina-associated domains (LADs; heterochromatin at the nuclear periphery) usually have a size between 10^4^ and 10^7^ bp (Steensel and Belmont, 2017).

A comparison with ChromHMM, Segway and EpiC-Seg suggests that EpiSegMix has the potential to provide segmentations that better reflect genomic annotations and yields states that are more predictive of gene expression. Moreover, the flexible duration modeling allows to effectively capture the reflective state lengths of long gene bodies and heterochromatic domains. The influence of the modified HMM topology on the segmentation suggests that testing other topology models is an important aspect to increase the modeling accuracy. Another direction of future work could be to combine the flexible duration modeling of EpiSegMix with a hierarchical HMM, as proposed by Marco et al. (2017), which may prove to be a powerful idea to perform interdependent chromatin segmentation at different length scales.

Although our results suggest that flexible distributions such as the Sichel or Beta Negative Binomial distribution often give the best results, we support a variety of distributions to deal with changing data properties and provide an easily extendable framework to integrate other data, such as ATAC-seq, DNase-seq or transcription factor ChIP-seq data.

In summary, we show that the modeling assumptions of the HMM have an impact on the segmentation quality and biological interpretation. Due to its high flexibility, EpiSegMix accurately fits HMM read count data with varying distributional properties and provides the additional option of flexible duration modeling. Finally EpiSegMix provides a widely configurable framework for chromatin segmentation that can be applied to a wide range of data.

## Declarations

## Author contributions

J.S. and N.A. contributed equally to this work. The method was developed by J.S. and N.A. under the supervision of J.W., A.S. and S.R.. L.L. performed preliminary analyses to select suitable distributions. J.S, N.A., J.W. and S.R. contributed to writing, and all authors contributed to revising the manuscript.

## Ethics declaration

None declared.

## Data availability

The data can be downloaded from the ENCODE portal. All accession numbers are provided in Supplementary Section S9.

## Acknowledgments

We acknowledge the ENCODE consortium for producing the data.

## Supplement

### S1. Supplementary Methods

#### S1.1. Notation for Hidden Markov Models (HMMs)

- finite set of hidden states {1, 2, …, *N* }
- emission alphabet Σ
- observation *O* = (*O*_1_, …, *O*_*T*_) with *T* time points, where *T* is the number of rows in the count matrix
- hidden state sequence *Q* = (*Q*_1_, …, *Q*_*T*_)
- probability vector (*π*_*i*_)_*i*∈{1,2,…,*N*}_, which defines the starting probabilities for all states
- *N* × *N* matrix *A*, where each entry *a*_*ij*_ denotes the transition probability to move from state *i* to state *j*
- state-specific emission parameters *B* = {*b*_*j*_(·) | *j* ∈ {1, 2, …, *N* }}
- probability distribution *P*_*θ*_(·), governed by a set of parameters *θ*

#### S1.2. Parameter Estimation (Baum-Welch Algorithm)

The parameter estimation for an HMM with parameters *θ* = (*π, A, B*) is performed using the Baum-Welch algorithm (Rabiner, 1989), which is an expectation-maximization algorithm tailored to HMMs. In each iteration *r*, the complete log likelihood function is maximized until convergence to a local optimum or until the maximum number of iterations has been reached. In our flexible distribution HMM, the complete log-likelihood function is given by Equation (6)

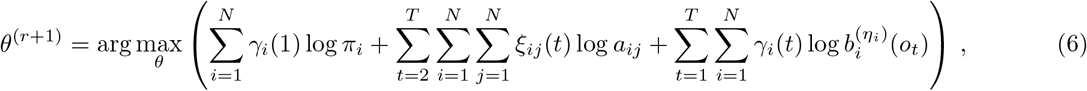

where the emission probabilities 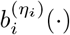are specified by the distribution type and its state-specific parameters *η*_*i*_ and

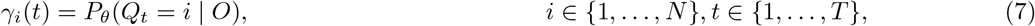

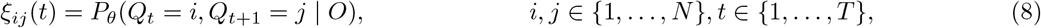

which are efficiently computed using the forward-backward algorithm (Rabiner, 1989), resulting in the following update rules (Bilmes, 1998):

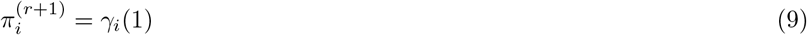

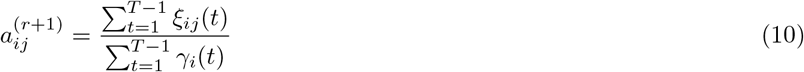

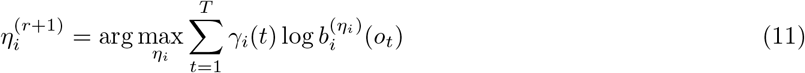

##### Multivariate observations (histone marks)

For an observation with *M* histone marks, the update formula for the emission parameters in state *i* is given by

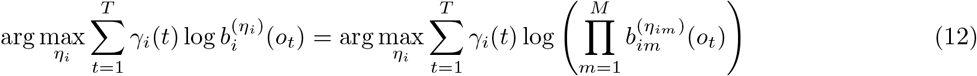

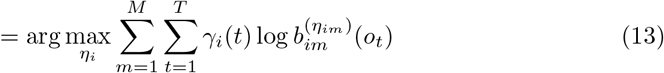

Thus, the parameters *η* can be optimized independently for each state *i* and each histone mark *m*:

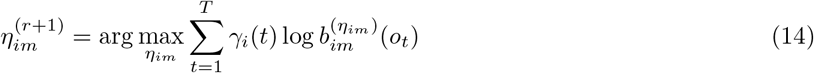

##### Numerical Maximum-likelihood optimization

To guarantee the convergence of the Baum-Welch algorithm to a local optimum, the maximum likelihood estimators are used to update the parameters of all distributions. Since for most distributions no closed-form solution exists for Equation (14), it is optimized numerically using the *MIGRAD()* function from the *ROOT Minuit 2* package (Brun and Rademakers, 1997). All computations are performed with logarithms of probabilities for numerical stability.

##### Extended-state HMM

In the extended-state HMM, for each state *i*, we create a sub-HMM with states *i*_1_, *i*_2_, …, *i*_*S i*_, where *S*_*i*_ is the number of states in the sub-HMM of state *i*. The update formulas enforcing the additional constraints of equal emission and self-transition probabilities are given by

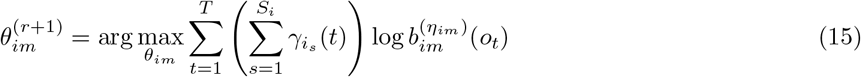

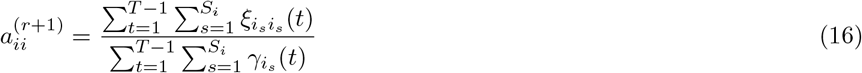

All other parts of the Baum–Welch algorithm remain unchanged.

##### Initialization

As an iterative technique, the Baum-Welch algorithm is sensitive to the given initial parameters. Therefore, systematically guessed initial values are beneficial both for convergence speed and goodness-of-fit. To find proper starting values, we first perform *k*-means clustering with *k* equal to the desired number of states. In the next step, the method-of-moments estimators are calculated for each cluster and used to initialize the emission parameters. The transition probabilities are initialized uniformly.

#### S1.3. Details on Available Emission Distributions

This section lists technical details about the available distributions for modeling the read counts (emissions of the HMM). We provide the probability mass function, the method-of-moments estimators for initialization and, if available, the closed form of maximum likelihood estimators (MLEs) for all implemented discrete distributions.

For the method-of-moments estimators, the sample mean 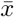 and sample variance *s*^2^ for a sample (*x*_1_, …, *x*_*N*_) are given by

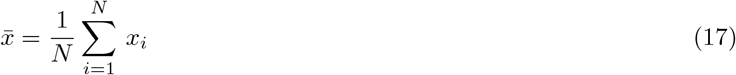

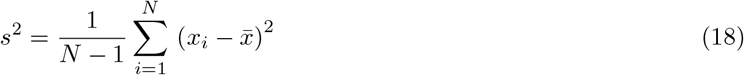

##### S1.3.1. Binomial Distribution

The probability mass function of a Binomial distribution (BI) with parameters 0 ≤*p* ≤ 1 and a natural number *n >* 0 is given by (Bernoulli, 1713)

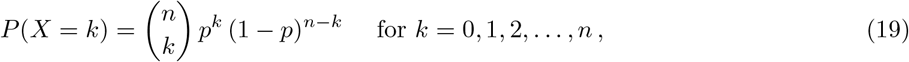

where 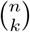 is the Binomial coefficient defined as 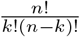. Assuming that *n* is known, the method-of-moments estimator of *p* is given by

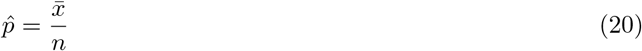

The MLE of *p* in state *i* is given for an observation *O* = (*o*_1_, …, *o*_*t*_) and membership coefficients *γ*_*i*_(*t*) by

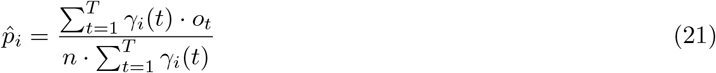

If *n* and *p* are unknown, the estimator of *n* is not guaranteed to be finite. Thus, we use the sample maximum as an estimator of *n*, as proposed by Fisher (1941).

##### S1.3.2. Poisson Distribution

The probability mass function of a Poisson distribution (PO) with parameter *λ >* 0 is given by (Poisson, 1837)

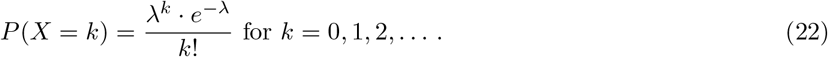

The method-of-moments estimator for *λ* is given by the sample mean (Johnson et al., 1993, ch. 4), and the MLE is computed for an observation *O* = (*o*_1_, …, *o*_*t*_) and membership coefficients *γ*_*i*_(*t*) by

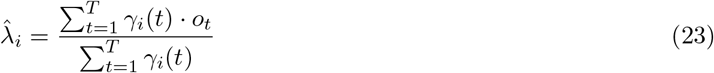

##### S1.3.3. Negative Binomial Distribution

The probability mass function of a Negative Binomial distribution (NBI) with parameters *r* ≥0 and 0 ≤*p* ≤ 1 is given by

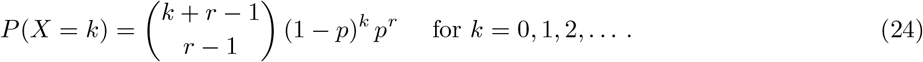

The method-of-moments estimators are given by (Lindén and Mäntyniemi, 2011)

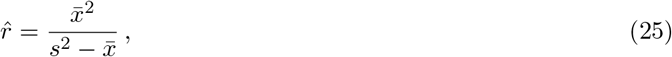

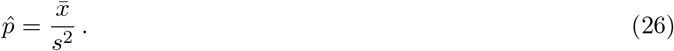

##### S1.3.4. Beta Binomial Distribution

The probability mass function of a Beta Binomial distribution (BB) with parameters *α >* 0, *β >* 0 and a natural number *n >* 0 is given by (Griffiths, 1973)

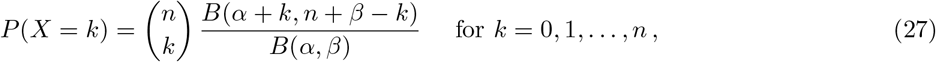

where

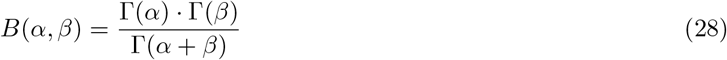

is the Beta function and

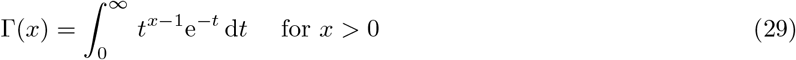

is the the Gamma function (Johnson et al., 1993, p.4). Other names for the Beta Binomial distribution are Pólya or Negative Hypergeometric distribution. The method-of-moments estimators of *α* and *β* are given by

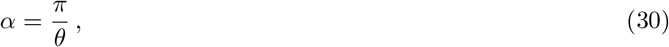

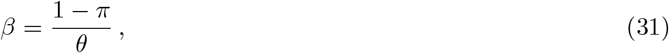

where *π* and *θ* are estimated by

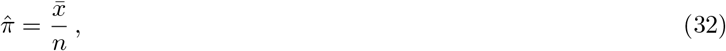

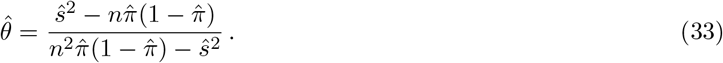

Similar to the Binomial distribution, if *n* is unknown, it is estimated by the sample maximum.

##### S1.3.5. Beta Negative Binomial Distribution

The probability mass function of a Beta Negative Binomial distribution (BNB) with parameters *r >* 0, *α >* 0, *β >* 0 is given by (Johnson et al., 1993, ch. 6.2.3)

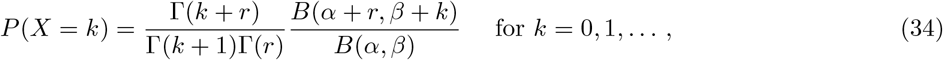

where *B*(·,·) denotes the Beta function (28) and Γ(·) denotes the Gamma function (29). Other names for the Beta Negative Binomial distribution are generalized Waring distribution, inverse Markov–Pólya distribution or Beta–Pascal distribution.

Since the *n*–th theoretical moment is only finite if *α > n* and in addition *r* and *β* are interchangeable (Rodríguez-Avi et al., 2007), a simplified approach is used to calculate the method-of-moment estimators. First, *r* and *p* are estimated under the assumption of a Negative Binomial distribution. The method-of-moments estimators of *α* and *β* are then estimated for a Beta distribution with the sample mean given by *p* and the sample variance given by an arbitrarily chosen value of 0.05. For 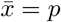 and *s*^2^ = 0.05, *α* and *β* are then given by (Schröder and Rahmann, 2017)

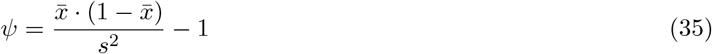

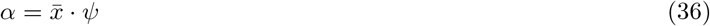

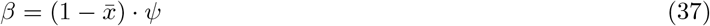

Since the method-of-moments estimators are only used for initialization of the Baum-Welch algorithm, inaccurate estimates are acceptable.

##### S1.3.6. Sichel Distribution

The probability mass function of a Sichel distribution with parameters *µ >* 0, *Σ >* 0 and −∞*< v <*∞ is given by (Sichel, 1992)

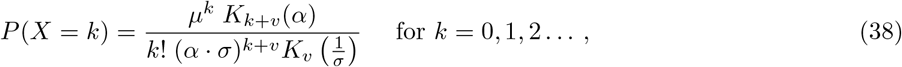

Where 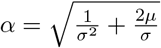 and *K* (*t*) denotes the modified Bessel function of the second kind of order *v* and argument *t* given by

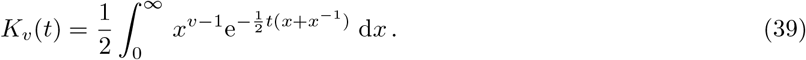

To avoid numerical problems during the computation of the Bessel function for large orders, the probability mass function is calculated recurrently based on the following recurrence relation (Stein et al., 1987)

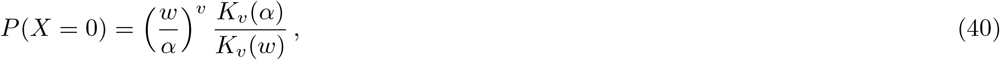

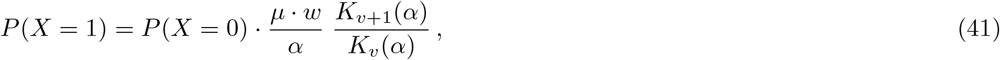

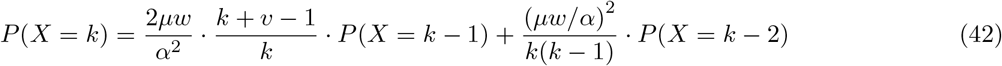

with 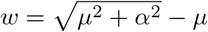.

For 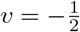 the Sichel distribution is equal to the inverse Gaussian Poisson distribution (Stein et al., 1987). For initialization, the parameters of the Sichel distribution are estimated using the method-of-moments estimators of the inverse Gaussian Poisson distribution, given by

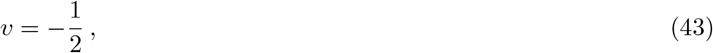

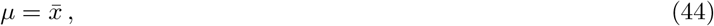

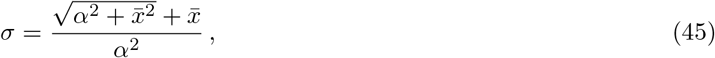

with 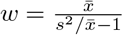 and *α*^2^ = (*w* + *µ*)^2^ − *µ*^2^.

##### S1.3.7. Zero-Adjusted Distributions

A zero-adjusted distribution is a discrete mixture distribution that has a fixed zero probability *π*. We currently support the zero-adjusted Poisson, Negative Binomial, Beta Negative Binomial and Sichel distribution. For a discrete random variable *X*^′^, the zero adjusted probability mass function with zero probability 0 ≤*π* ≤ 1 is given by (Rigby et al., 2019)

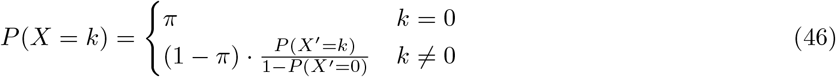

The method-of-moments estimator of *π* is given by the zero frequency and the MLE of *π* is given by

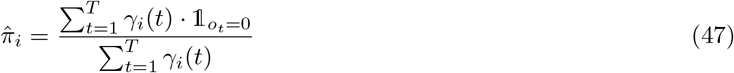

where *γ*_*i*_(*t*) denotes the membership coefficient of observation *o*_*t*_ in state *i*. The parameters of the distribution for *X*^′^ can be optimized independently.

### S2. Selecting the Number of States

Since the log-likelihood of an HMM increases monotonically as the number of states increases, the selected number of states should be a good compromise between model fit and interpretability. Figure 7 shows the log-likelihood (LL) curve for different numbers of states on one of the cell lines. While the LL increases steeply for small numbers of states, it flattens for larger numbers; so 10 states emerges as a good compromise between goodness-of-fit and biological interpretability.

**Figure 7.**
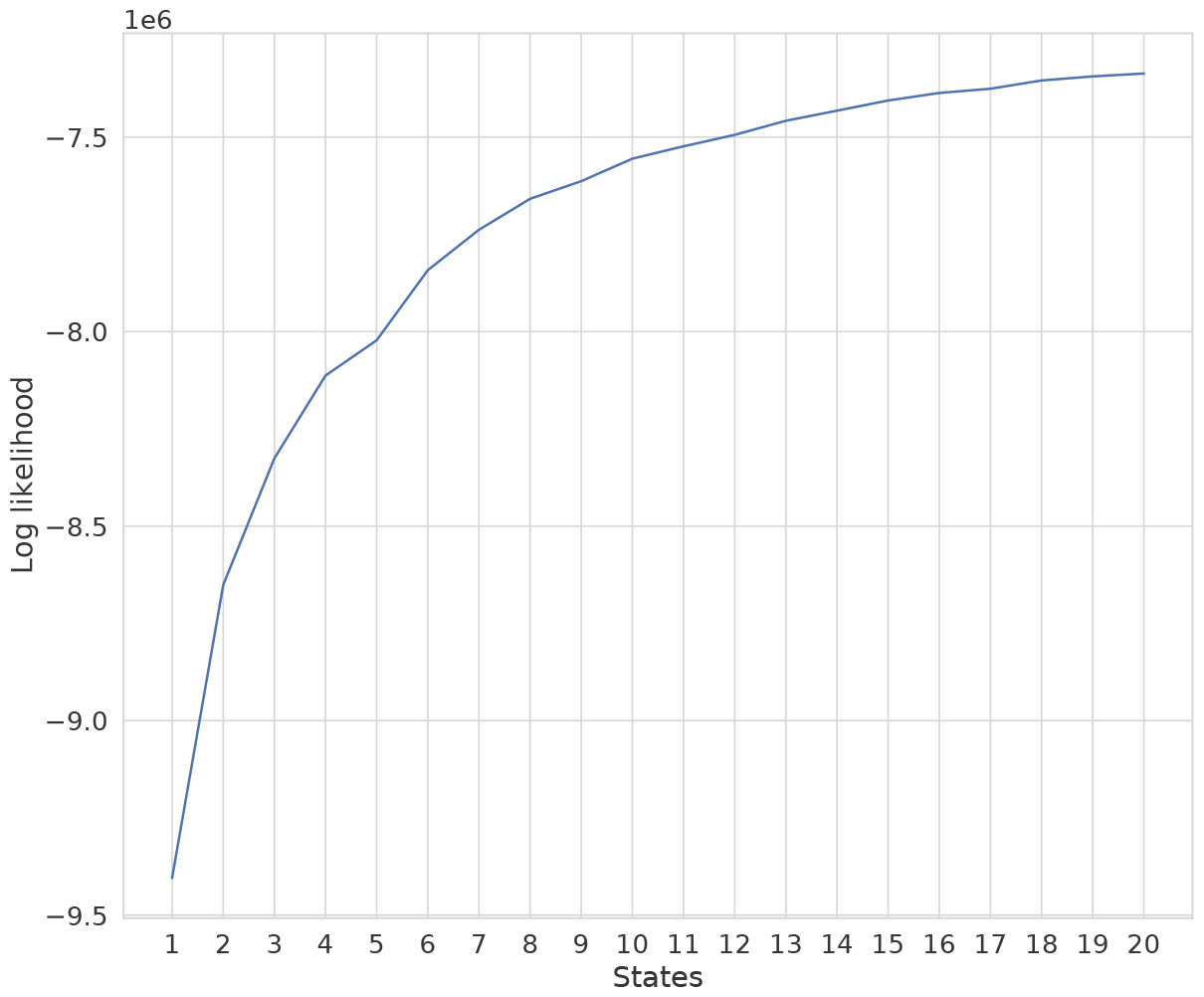
Log-likelihoods for a 1-state HMM up to a 20-state HMM for the IMR90 1 dataset using the six core histone marks H3K9me3, H3K27me3, H3K36me3, H3K4me1, H3K4me3 and H3K27ac. A higher log likelihood indicates a better model fit.

### S3. Data Quality

*Deeptools* and *multiqc* were used to validate the quality of the data. All alignment files in BAM format were initially processed using *deeptools*. A multibam report was generated using *multiqc*. Figure 8 depicts the fingerprint curves of the genomic distribution of read counts of histone marks across various samples. A diagonal line would indicate a uniform enrichment of a histone mark across the genome (no selective enrichment). Broad marks such as H3K9me3 and H3K27me3 are expected to have curves near the diagonal line, since they are enriched in a large part of the genome. Conversely, narrow marks such as H3K27ac show a sharp curve called elbow point, where the peaks cover a small fraction of the genome. Figure 8 demonstrates that the data quality and amount of noise varies across the six cell lines and their replicates. Hence, our evaluation covers a broad spectrum of old and recent data of variable quality.

**Figure 8.**
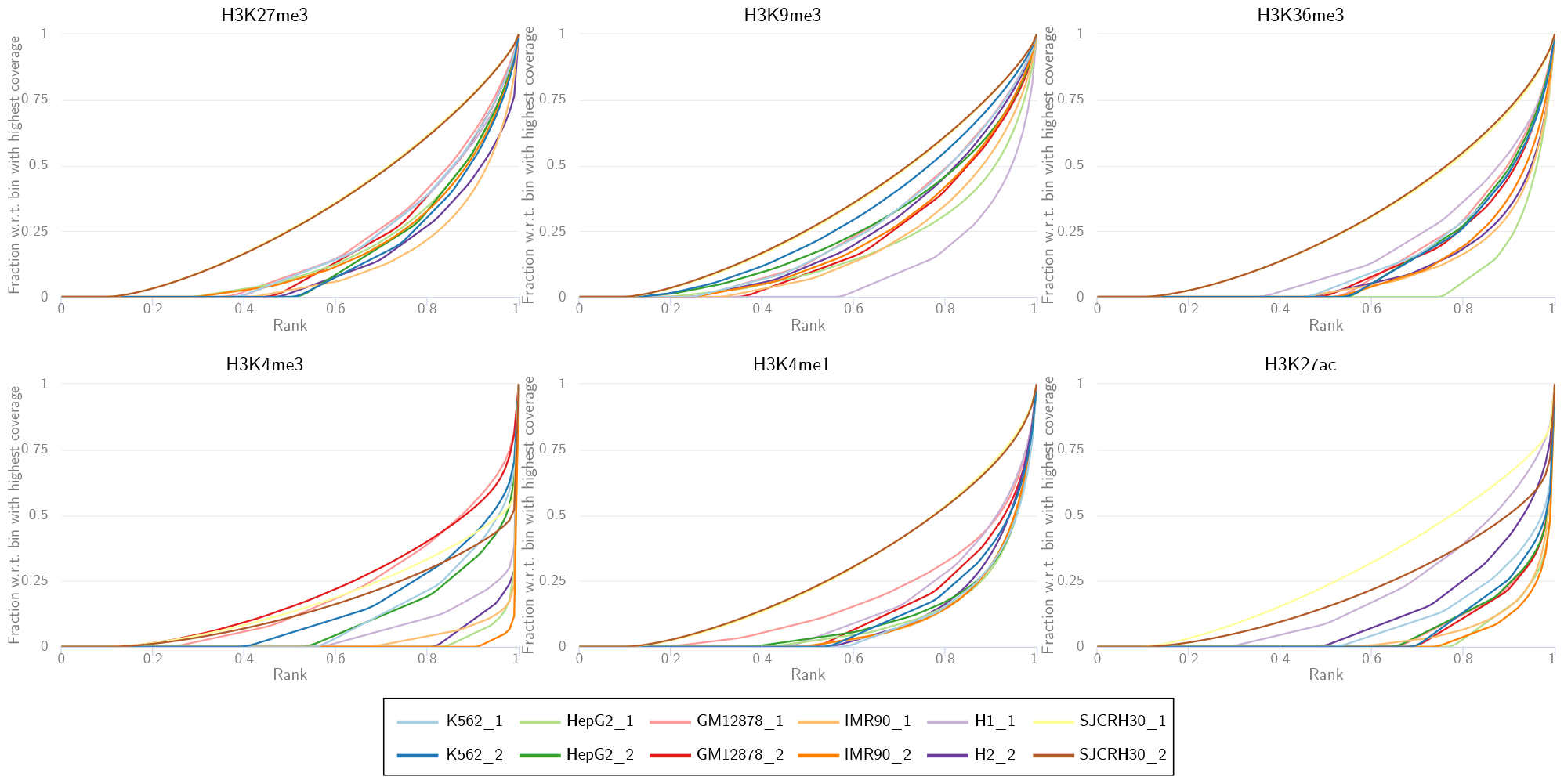
Quality of datasets by evaluating the genome-wide ChIP-seq enrichment for all histone marks (panels) in the different cell lines (colors). The curves show the fraction of reads (y-axis) needed to cover a certain fraction of the genome (x-axis). A diagonal line would indicate uniform enrichment of the histone mark across the genome (noise, input, no signal). A curve close to the x-axis and the vertical line at *x* = 1 indicates a highly selectively enriched histone mark. The fingerprint plots were created using *deeptools* (Fidel et al., 2016) *and multiqc* (Ewels et al., 2016).

### S4. Flexible Emission Modeling

#### S4.1. Selecting the Most Appropriate Distributions

To determine which distribution is best suited to model the read counts for each histone modification, we provide a workflow that tests, for each histone modification, a set of distributions and outputs a model file that can be used for the main segmentation workflow (https://gitlab.com/rahmannlab/episegmix). For each mark, a 3-state HMM is fitted individually for the different distributions. We then choose the distribution with the highest log-likelihood. Table 2 lists the used distribution for the results shown in the main article.

**Table 2:** Selected distribution for each mark per dataset. NBI denotes the Negative Binomial, SI the Sichel and BNB the Beta Negative Binomial distribution. For cell lines SJCRH30 1 and GM12878 1, the Sichel distribution was used for all marks due to a better quality in the combined model.

| cell line | histone mark |  |  |  |  |  |
| --- | --- | --- | --- | --- | --- | --- |
|  | H3K9me3 | H3K27me3 | H3K36me3 | H3K4me1 | H3K4me3 | H3K27ac |
| K561_1 | SI | SI | SI | SI | SI | SI |
| K562_2 | BNB | NBI | BNB | SI | BNB | SI |
| HepG2_1 | BNB | BNB | BNB | SI | BNB | SI |
| HepG2_2 | BNB | BNB | NBI | SI | SI | SI |
| H1_1 | SI | BNB | BNB | SI | BNB | SI |
| H1_2 | SI | BNB | SI | SI | SI | SI |
| SJCRH30_1 | SI | SI | SI | SI | SI | SI |
| SJCRH30_2 | SI | SI | SI | NBI | SI | BNB |
| IMR90_1 | BNB | NBI | SI | SI | SI | SI |
| IMR90_2 | BNB | NBI | BNB | SI | SI | SI |
| GM12878_1 | SI | SI | SI | SI | SI | SI |
| GM12878_2 | BNB | BNB | NBI | SI | BNB | SI |

In most cases, the flexible three-parametric distributions (Sichel and Beta Negative Binomial distribution) were selected. In addition, the same distribution is not consistently chosen for the same histone mark which can be explained by the previously discussed differences in the data quality (see Supplementary Section S3).

#### S4.2. Impact of Distribution Types

Figure 9 and Figure 10 show the advantage of using flexible emission modeling. Comparing the Log-Likelihood value between models with negative binomial and flexible emissions shows that the model fit of the flexible model is consistently better on all tested datasets. In addition, Figure 10 highlights that a theoretical better model fit also impacts the biological interpretability. In particular, narrow marks, such as H3K4me3, H3K4me1 and H3K27ac, can more accurately be modeled using flexible distributions, such as the Sichel distribution. For example, the Negative Binomial distribution cannot model the highly skewed read count distribution of H3K4me3 in state 3, where the high variance can only be modeled by selecting a parameter combination that additionally leads to a high probability for small read counts. In contrast, the Sichel distribution (as selected by the previously applied distribution fitting), is able to model the skewed and highly variable read counts of H3K4me3 without the above mentioned limitation. This also reflects in a lower genomic coverage of state 3. Since H3K4me3 is typically enriched in promoters while H3K4me1 is enriched in active enhancers, the results suggest that flexible distributions can better distinguish between promoters and enhancers.

**Figure 9.**
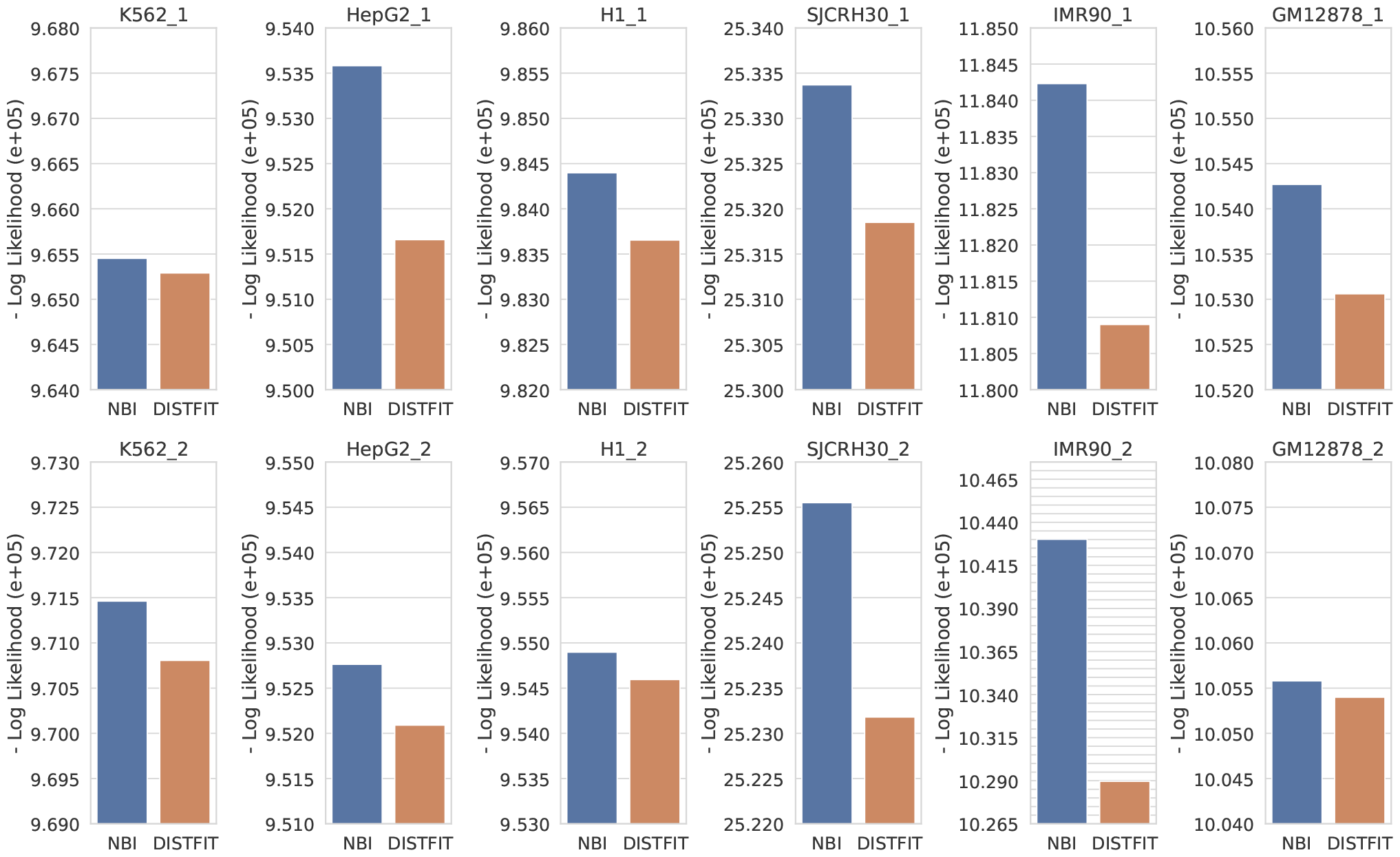
Comparison of the negative Log-Likelihood values between HMMs using Negative Binomial and flexible emissions. A lower negative Log-Likelihood value indicates a better model fit. For the *DISTFIT* model, the distributions were selected according to Table 2. Since the Log-Likelihood is only comparable across the same dataset, the plot is split by dataset. Note that the y-scale does not start at 0 to highlight the differences. Each grid line corresponds to an increase of 0.005 · 10^5^.

**Figure 10.**
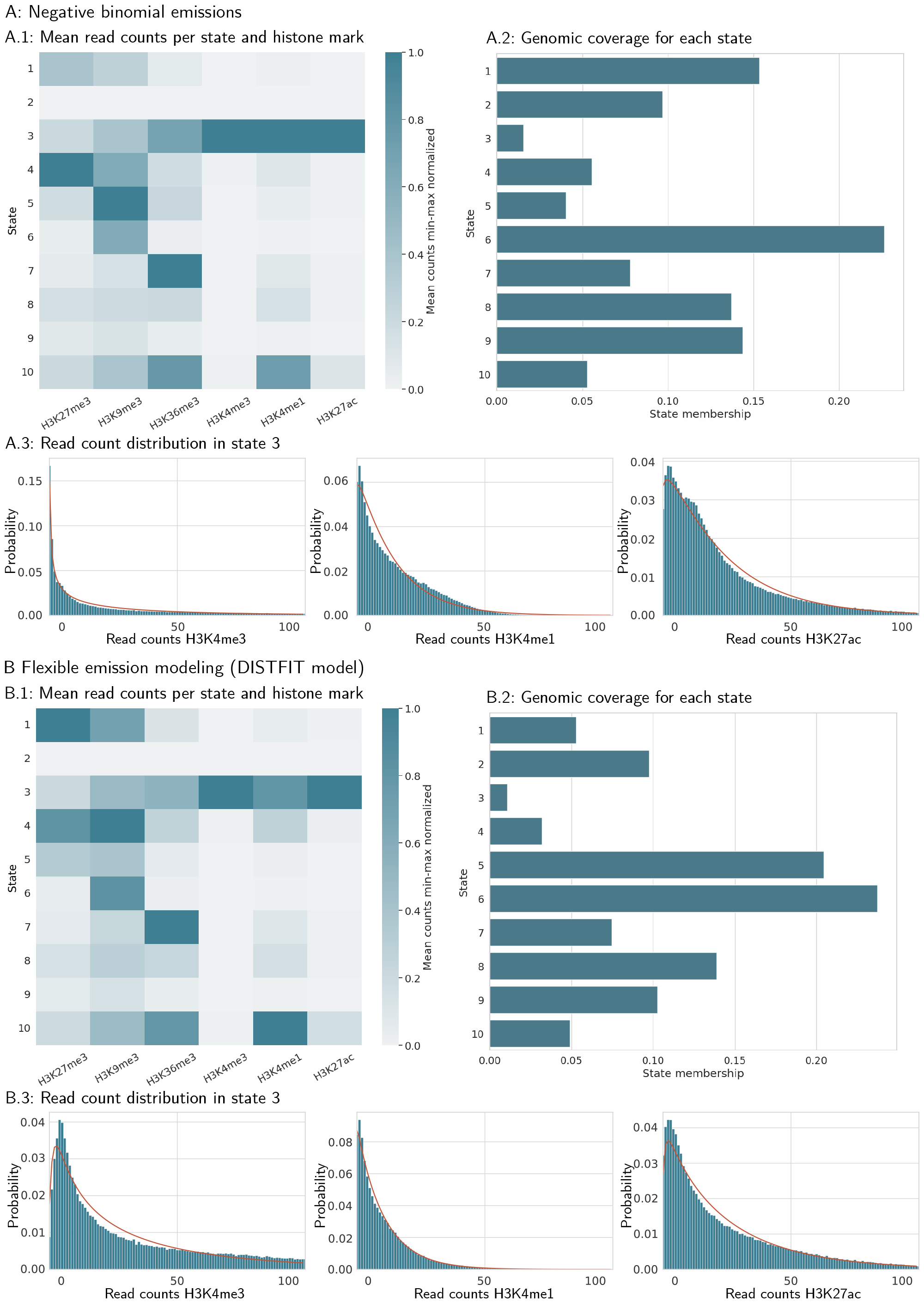
Effects of emission modeling on the discovery of chromatin states illustrated by IMR90 2 (**A**: negative binomial emissions vs. **B**: flexible emission modeling according to Table 2). In particular, the comparison of state 3 and 10 highlights the impact of choosing different distribution types.

### S5. Impact of Duration Modeling

Figure 11 shows the segment length distribution (consecutive 200 bp bin assigned to the same state) and the normalized histone enrichment per state for an HMM with a classic topology (Geometric state duration) and an extended state HMM (Negative Binomial state duration). Both models show similar histone mark patterns but differing segment lengths. For example, the segment lengths of state 3 (heterochromatin, H3K27me3 and H3K9me3) and state 5 (transcriptional elongation, H3K36me3) are longer for the extended-state HMM in comparison to the classic HMM topology. Since the histone marks associated with active genes and heterochromatin are usually enriched as broad domains and not narrow peaks, we may presume that flexible duration modeling allows to more accurately model the length of biological domains.

**Figure 11.**
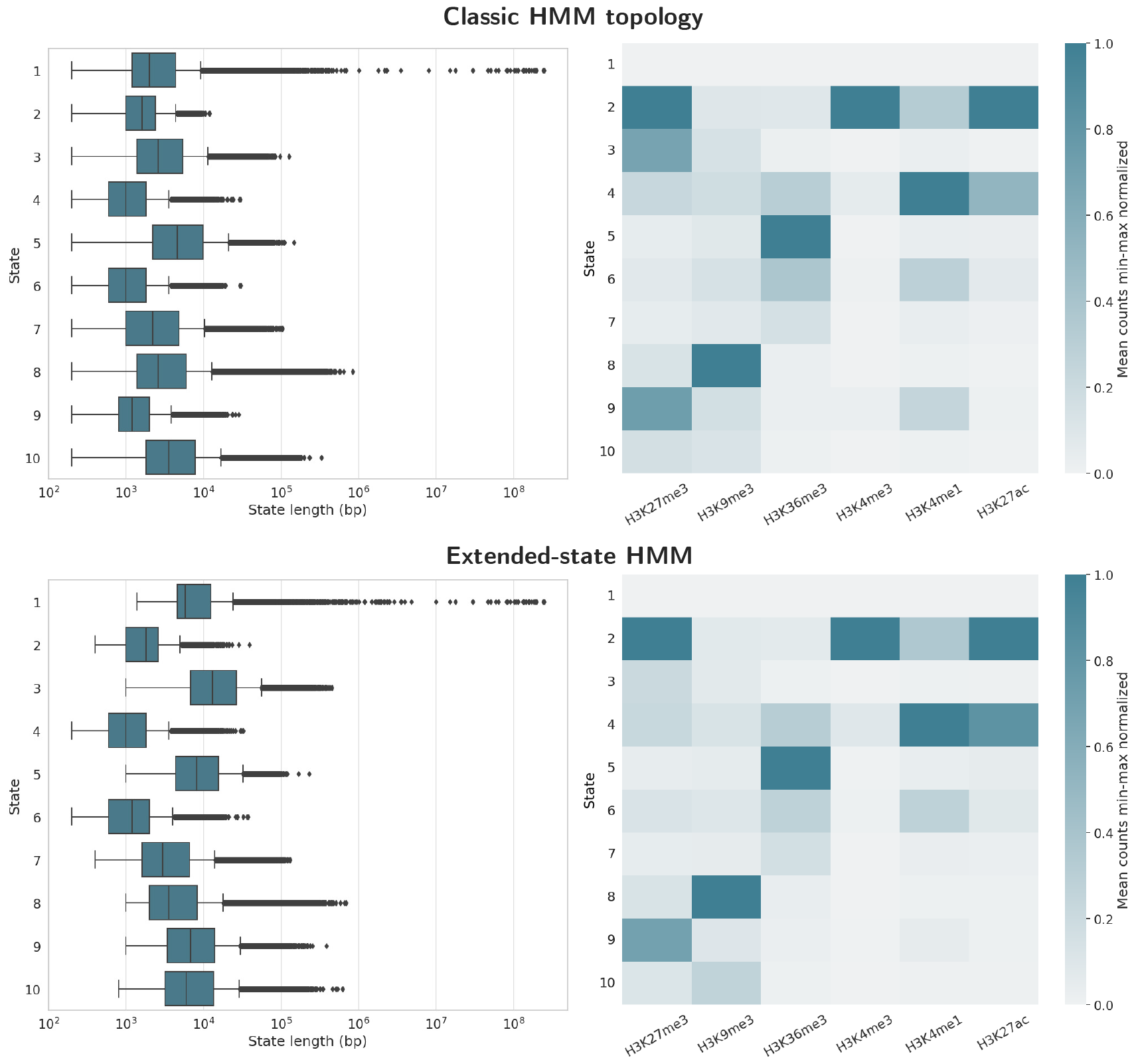
Comparison of segmentations trained with an HMM with classic topology and a flexible duration HMM (extended-state topology) for HepG2 1 using the emission distributions given in Table 2.

### S6. Robustness

To evaluate the robustness of the different chromatin segmentation methods, we generated the count matrix for two technical replicate experiments each. However, evaluating the robustness of chromatin segmentation methods depends on many factors that make general statements difficult. For example, a higher number of states usually results in a lower robustness due to interchangeable states. In addition, robustness often differs between active and repressive states (usually lower for regulatory states), which requires to evaluate the robustness after biological interpretation. Since so far no gold standard for automatic annotation exists, this approach may be flawed by manual or inaccurate automatic annotation. Hence, we compare the robustness using three different measures that capture different properties a robust method should fulfill.

First, we compared the Jaccard index, averaged over all states, between the segmentations of the two replicates after mapping the states of both segmentations according to the highest Jaccard index. Given a set of bins *b*_1_ that are assigned to the state in the first replicate and a set of bins *b*_2_ assigned to the corresponding state in the second replicate, the Jaccard index is defined as the size of the intersection *b*_1_ ∩ *b*_2_ divided by the size of their union *b*_1_ ∪ *b*_2_, and hence is between 0 (no agreement at all) and 1 (complete agreement). A higher index reflects higher robustness. Since this approach does not require a prior biological annotation, it is independent of potential biases during annotation. Figure 12 shows that the average robustness of EpiSegMix is slightly higher compared to the other methods which may be due to a high robustness in the heterochromatic states.

**Figure 12.**
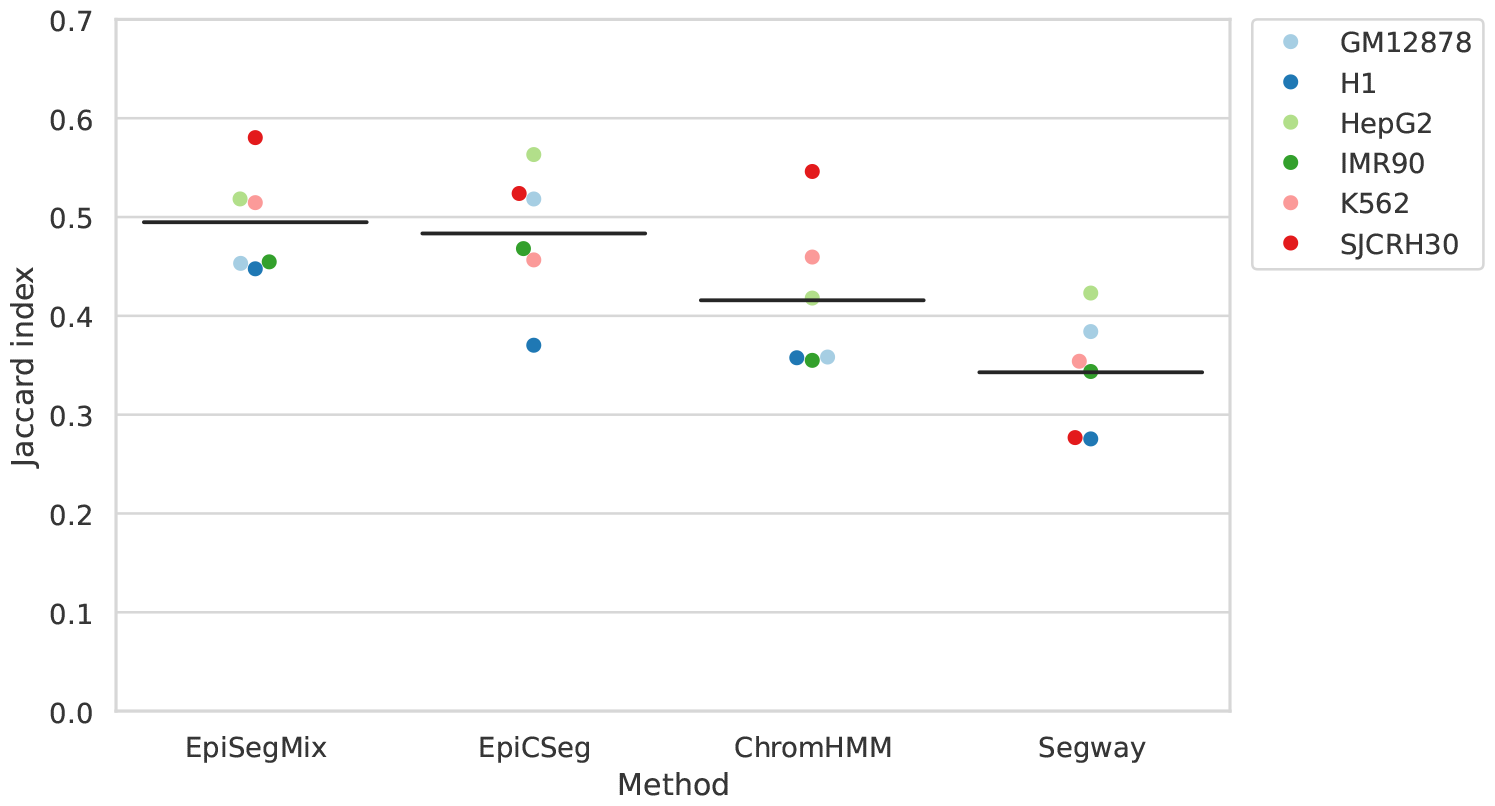
Robustness between replicate datasets. Each point represents the average Jaccard similarity between the segmentations of two technical replicates, with a higher Jaccard index indicating a greater overlap.

For a more detailed evaluation of the robustness we first manually annotated each state with one of 10 biological categories. Figure 13A shows the percentage of how many bases in the first replicate are covered by the closest matching state in the second replicate. Thus, if there existed a perfect one-to-one correspondence between both replicates each overlap would be 100%. In general, the robustness of all methods strongly varies between cell lines and biological annotations, which suggests that the robustness of all methods strongly depends on the data quality. EpiSegMix shows a higher robustness for promoter, transcription and strong heterochromatic states, but has a lower robustness for enhancers. However, due to the high variation we refrain from drawing a general conclusion.

**Figure 13.**
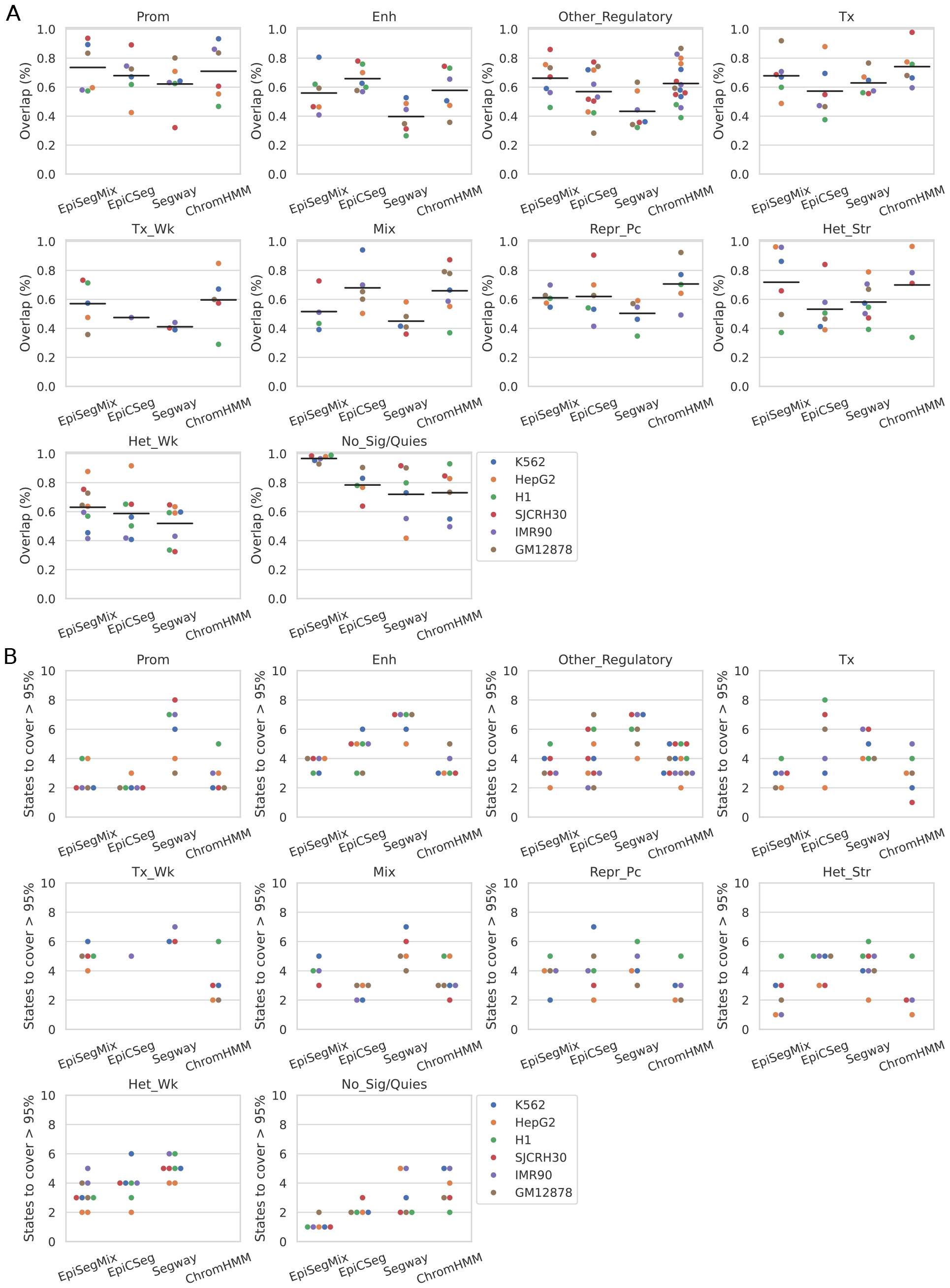
Robustness per state (each state in the first replicate was manually annotated). **A** The percentage of how many base pairs are covered by each state in the second replicate was calculated. Each dot corresponds to the percentage covered by the state with the highest overlap. **B** Number of states of the second replicate that are required such that over 95% of the state in the first replicate are covered.

Since a lower overlap may be caused by several states with very similar properties, we further calculated how many different states in the segmentation of the second replicate are required to cover over 95% of each state in the segmentation of the first replicate. Figure 13B shows the number of states for each annotated state, where a lower number of states indicates a higher reproducibility of finding states with a similar biological interpretation. In comparison to the other methods, EpiSegMix usually requires a similar or lower number of states to cover 95% of each state.

In conclusion, all methods have a limited robustness to replicate datasets, which may be caused by noise or a too fine-grained segmentation. In all considered measures, EpiSegMix had a similar (or sometimes better) robustness compared to other methods.

### S7. Supplementary Figure K562

The following 3 figures provide an in-depth look at the results for cell line K562. Figure 14A displays two genomic regions of distinct transcription activity; Figure 14B shows the enrichment patterns of each histone mark across HMM states for each method. Figure 15 visualizes state enhancement near validated enhancers. Figure 16 shows gene activity in regions that ChromHMM classified as weak transcription (Tx Wk; panel A) or quiescent (Quies; panel B), grouped by the predicted EpiSegMix state. EpiSegMix estimates slightly different acitivity levels per state.

**Figure 14.**
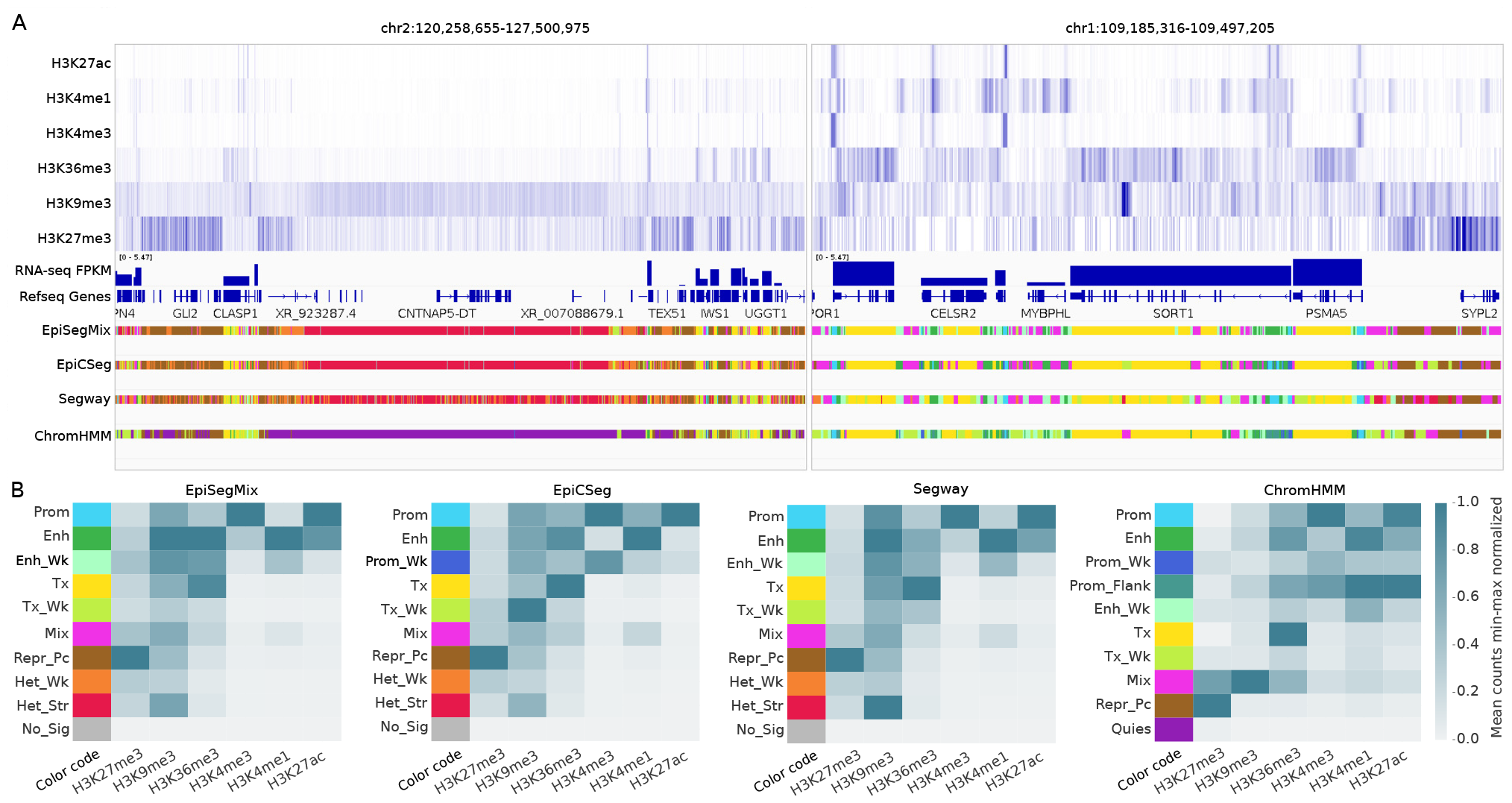
IGV-screenshot for two regions with different genomic activities. **A** The first six tracks show the ChIP-seq intensities of the six input histone marks. The 7th track shows the signal of unique reads from an RNA-seq experiment, and the 8-th track shows the Refseq gene positions. The last four tracks contain the segmentation of EpiSegMix, EpiCSeg, Segway and ChromHMM. On the left, a heterochromatic region is shown which highlights the smooth transition from strong heterochromatin (high H3K9me3) to Polycomb repressed heterochromatin (high H3K27me3). On the right, an actively transcribed region is shown. **B** Heatmap showing the histone patterns in the discovered chromatin states with color legend.

**Figure 15.**
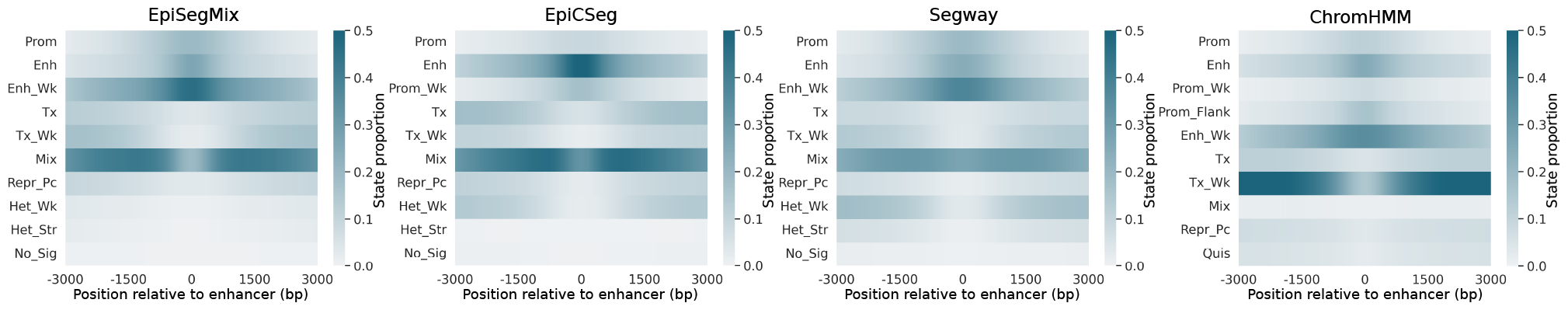
State enrichment near enhancers in K562. Validated enhancers were downloaded from http://enhanceratlas.org/ (Gao et al., 2016) and the state annotation 3000 bp up- and downstream was evaluated. The heatmap shows for each position the percentage of each state.

**Figure 16.**
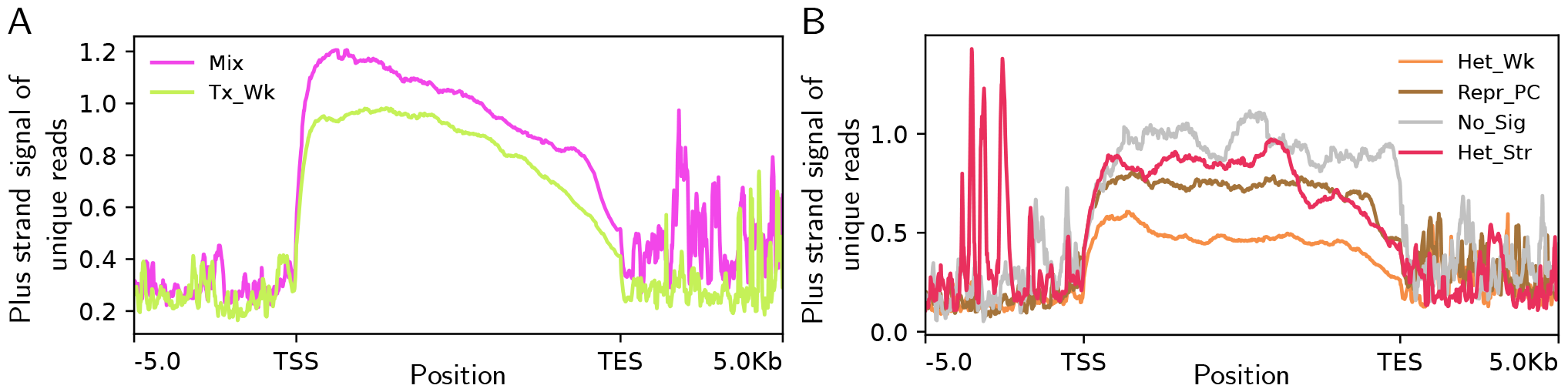
ChromHMM reclassified genes in EpiSegMix. The figure shows the reallocated genes of ChromHMM’s Quies state and Tx Wk states to EpiSegMix well defined states that reflect distinct gene activity. Each gene was annotated with maximum overlapping (in base pairs) states across different methods. All genes were stretched or squished to 10 Kbp along 5 Kbp up and downstream. **A** The protein coding genes that overlapped with Tx Wk state in ChromHMM and Mix or T Wk in EpiSegMix were used to plot the gene activity across and around genes using *deeptools* (Fidel et al., 2016). **B** Similar to Figure A, here showing the transition of genes from ChromHMM’s Quies to EpiSegMix heterochromatic states.

### S8. Biological Annotation of Chromatin States

We provide names, abbreviations and descriptions of the biological names (labels) of the HMM states.

| Name | Abbreviation | Description |
| --- | --- | --- |
| Promoter | Prom | Promoter state characterized by an enrichment of H3K4me3. Mainly found near transcription start sites. |
| Enhancer | Enh | Enhancer state showing an enrichment of H3K4me1 (and H3K36me3 in intergenic enhancers). |
| Enhancer Weak | Enh_Wk | Same as enhancer state but with lower signal intensities (often flanking enhancer state). |
| Promoter Weak | Prom_Wk | Similar enrichment to promoter state but with lower signal intensities. |
| Promoter Flanking | Prom_Flank | Low enrichment of promoter-associated marks and mainly flanking promoter regions. |
| Transcription | Tx | State labeling actively transcribed genes (enriched in gene bodies) and showing high signal intensities of the transcription elongation mark H3K36me3. |
| Transcription Weak | Tx_Wk | Similar to transcription state but with lower intensities of H3K36me3. |
| Mix | Mix | State showing an enrichment of several active marks (transcription, regulatory...) and a low intensity of repressive marks, found as intermediate/border state in active regions. |
| Repressive Polycomb | Repr_Pc | Polycomb repressed heterochromatin, characterized by high levels of H3K27me3. Important for cell-type specific gene repression. |
| Heterochromatin Strong | Het_Str | State showing an enrichment of the broad heterochromatic mark H3K9me3 (and H3K27me3). |
| Heterochromatin Weak | Het_Wk | Similar to strong heterochromatin, but with lower signal intensities. |
| Quiescent | Quies | Low signal intensity for any mark. |
| No Signal | No_Sig | No signal for any mark (mainly unmappable regions). |
| Regulatory | Other_Regulatory | Label for all states that show a mixed enrichment of regulatory marks (H3K27ac, H3K4me1, H3K4me3) and are not the main promoter or enhancer state with strongest enrichment of the respective histone modifications. |

### S9. Data Availability

We provide ENCODE accession numbers to all datasets used in the main article.

#### S9.1. Histone ChIP-seq Data

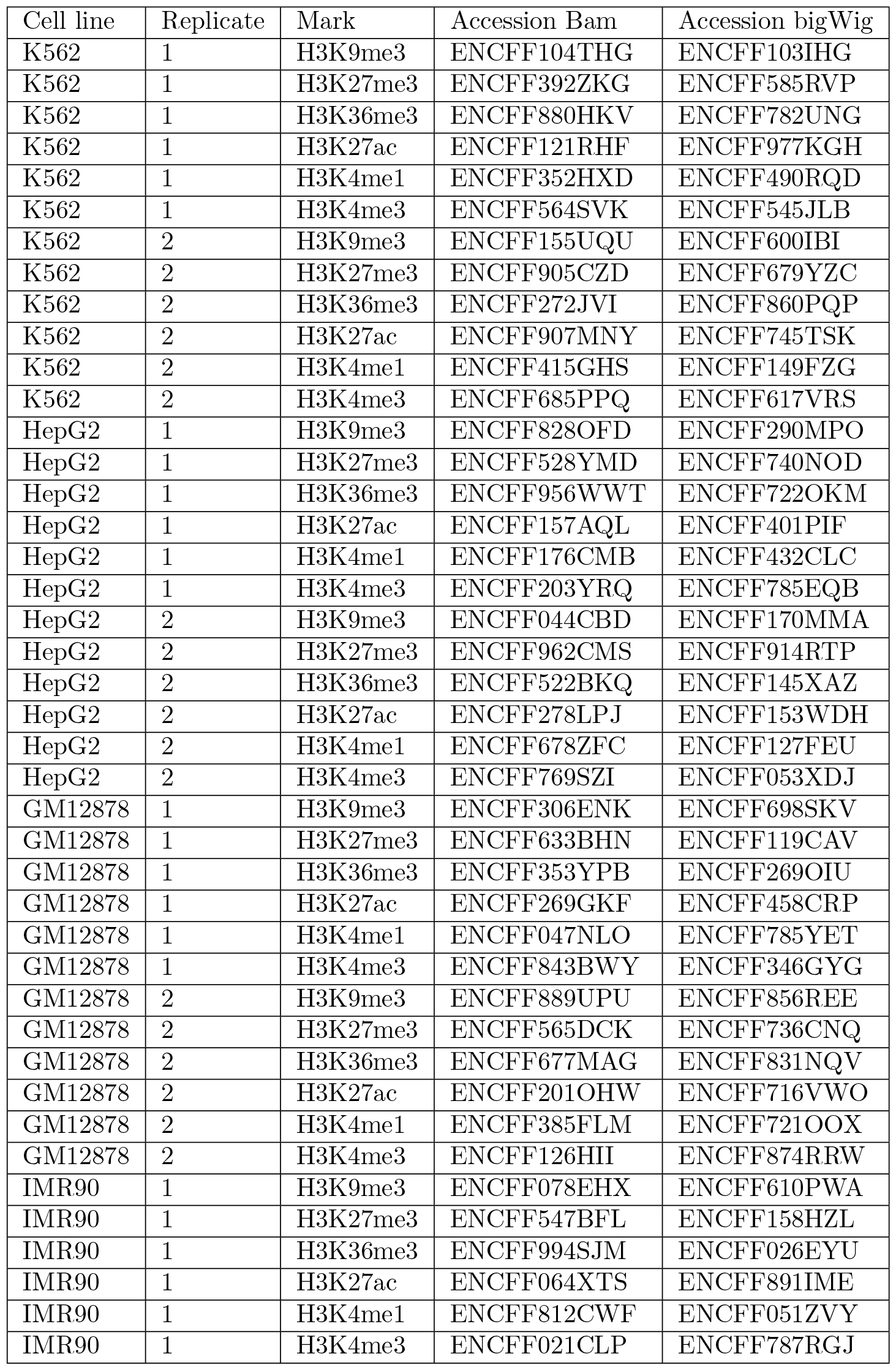

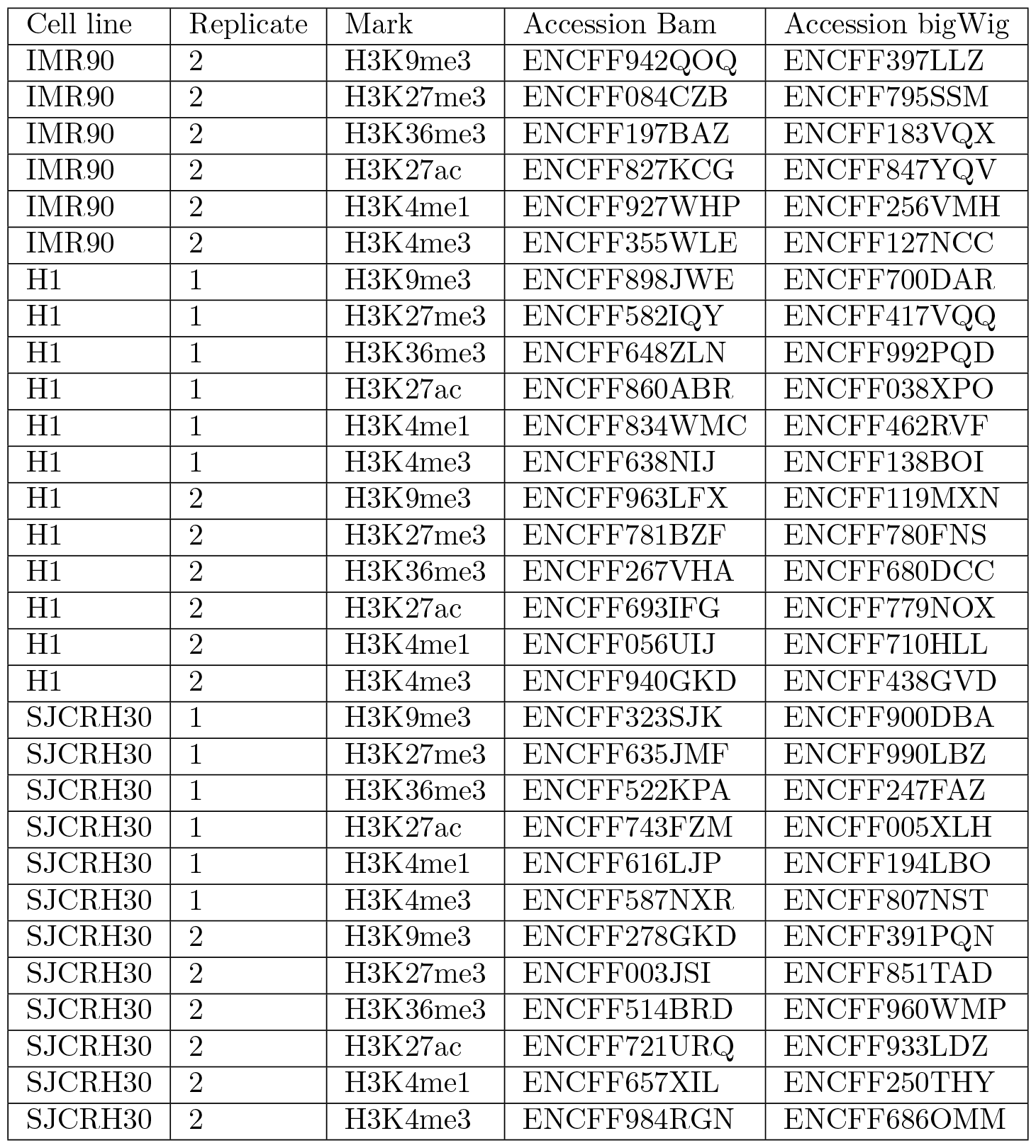

#### S9.2. RNA-seq Data

S9.2 RNA-seq Data
| Cell line | Replicate | Accession |
| --- | --- | --- |
| K562 | 1 | ENCFF068NRZ |
| K562 | 2 | ENCFF928YLB |
| HepG2 | 1 | ENCFF355TDA |
| HepG2 | 2 | ENCFF073RKC |
| GM12878 | 1 | ENCFF678BLG |
| GM12878 | 2 | ENCFF897XES |
| IMR90 | 1 | ENCFF958WFW |
| IMR90 | 2 | ENCFF631BAG |
| H1 | 1 | ENCFF243GKW |
| H1 | 2 | ENCFF432RPO |
| SJCRH30 | 1 | ENCFF771NMO |
| SJCRH30 | 2 | ENCFF955BPJ |

#### S9.3. ATAC-seq Data

S9.3 ATAC-seq Data
| Cell line | Accession |
| --- | --- |
| K562 | ENCFF512VEZ |
| HepG2 | ENCFF990VCP |
| GM12878 | ENCFF981FXV |
| IMR90 | ENCFF848XMR |

